# Estimating and accounting for genotyping errors in RAD-seq experiments

**DOI:** 10.1101/587428

**Authors:** Luisa Bresadola, Vivian Link, C. Alex Buerkle, Christian Lexer, Daniel Wegmann

## Abstract

In non-model organisms, evolutionary questions are frequently addressed using reduced representation sequencing techniques due to their low cost, ease of use, and because they do not require genomic resources such as a reference genome. However, evidence is accumulating that such techniques may be affected by specific biases, questioning the accuracy of obtained genotypes, and as a consequence, their usefulness in evolutionary studies. Here we introduce three strategies to estimate genotyping error rates from such data: through the comparison to high quality genotypes obtained with a different technique, from individual replicates, or from a population sample when assuming Hardy-Weinberg equilibrium. Applying these strategies to data obtained with Restriction site Associated DNA sequencing (RAD-seq), arguably the most popular reduced representation sequencing technique, revealed per-allele genotyping error rates that were much higher than sequencing error rates, particularly at heterozygous sites that were wrongly inferred as homozygous. As we exemplify through the inference of genome-wide and local ancestry of well characterized hybrids of two Eurasian poplar (*Populus*) species, such high error rates may lead to wrong biological conclusions. By properly accounting for these error rates in downstream analyses, either by incorporating genotyping errors directly or by recalibrating genotype likelihoods, we were nevertheless able to use the RAD-seq data to support biologically meaningful and robust inferences of ancestry among *Populus* hybrids. Based on these findings, we strongly recommend carefully assessing genotyping error rates in reduced representation sequencing experiments, and to properly account for these in downstream analyses, for instance using the tools presented here.

## Introduction

Despite the impressive advancements in sequencing techniques and the decrease of related costs, whole genome sequencing (WGS) remains prohibitively expensive when working with a large number of samples or species with large genomes. Since many applications do not require information on the whole genome, reduced representation sequencing techniques are valuable alternatives and have become widely used for genome-wide SNP discovery and genotyping, especially in species with poor genomic resources (Andrews, Good, Miller, Luikart, & Hohenlohe, 2016; Narum, Buerkle, Davey, Miller, & Hohenlohe, 2013).

A commonly used reduced representation sequencing technique is Restriction site Associated DNA sequencing (Baird et al., 2008; Miller, Dunham, Amores, Cresko, & Johnson, 2007), which allows the sequencing of massively multiplexed samples at minimal costs by focusing on the sequences adjacent to restriction sites. Since restriction sites are often shared between individuals within a species and often also between closely related species (Cariou, Duret, & Charlat, 2013), focusing on adjacent sequences guarantees that sequenced loci are mostly overlapping across samples.

Briefly, the first step in the original RAD-seq protocol is the digestion of genomic DNA with a restriction enzyme. The resulting fragments are ligated to an adaptor and a unique barcode for each sample, and multiple individuals are pooled. The fragments are then sheared using a sonicator and those showing the proper size are selected and amplified through polymerase chain reaction (PCR). At this point the library is suitable for sequencing. By focusing the sequencing effort on tagged restriction sites, rather than on all randomly sheared genomic fragments (Arnold, Corbett-Detig, Hartl, & Bomblies, 2013; Rowe, Renaut, & Guggisberg, 2011), the number of markers can be customized through the choice of restriction enzymes. This choice will also influence which features of the genome are sampled, since certain enzymes preferentially cut in exonic regions, while others target intergenic and intronic regions (Arnold et al., 2013; Pootakham et al., 2016).

Several alternative RAD-seq protocols allow for ample customization of this methodology. These include the elimination of the sonication step (Andrews et al., 2016), ddRAD (Peterson, Weber, Kay, Fisher, & Hoekstra, 2012), which uses two restriction enzymes rather than one, 2bRAD (Wang, Meyer, McKay, & Matz, 2012), which uses IIb-type restriction enzymes and produces fragments of 36 bp, and ezRAD (Toonen et al., 2013), in which DNA is digested with isoschizomers (a pair of restriction enzymes recognizing the same sequence). However, each method has different advantages and pitfalls, and a specific protocol may be more suitable for some applications than for others (Puritz et al., 2014). Due to this versatility, RAD-seq has been used in diverse applications, including the study of the genomics of adaptation (Andrews et al., 2016), hybridization and speciation (Marques et al., 2016), inbreeding depression (Hoffman et al., 2014), genetic associations (Nadeau et al., 2014), genetic mapping (Chutimanitsakun et al., 2011) and phylogeographic and phylogenomic analyses (Emerson et al., 2010; Leaché et al., 2015).

Despite this widespread use, genotypes called from RAD-seq data have been associated with several biases, many of which are specific to RAD-seq. Several major biases potentially affect alleles differently, which may lead to their unequal representation in sequencing data, and hence to genotyping errors at heterozygous sites (Davey et al., 2013). For instance, polymorphisms occurring in the restriction site may result in one allele not being cut and therefore not sequenced, potentially causing linked sites to be erroneously called homozygous (“allele dropout”). Polymorphisms at neighbouring restriction sites may also result in genotyping biases, for example, if the length of the fragment of one allele falls short of the selected size range (Andrews et al., 2016). Yet size differences among longer fragments were also found to result in unequal sequencing depth at linked sites because sonicators shear shorter fragments less efficiently than longer fragments (Sambrook & Russell, 2006). Finally, the PCR step present in most RAD-seq protocols may contribute to genotyping errors through unequal amplification of the two alleles (Casbon, Osborne, Brenner, & Lichtenstein, 2011) or through so-called PCR duplicates, the sequencing of misleading clonal copies of the same initial molecule (Andrews & Luikart, 2014, Euclid 2019). Since many protocols produce single-end libraries or libraries where both ends are defined by restriction sites (e.g. ddRAD), PCR duplicates cannot be reliably identified bioinformatically unless very many different adapter sequences are used (Schweyen, Rozenberg, & Leese, 2014). This is particularly problematic in the case of PCR errors that might be sequenced in many copies, resulting in wrongly called heterozygous genotypes (Andrews et al., 2016).

Some consequences of these biases in downstream analyses are well documented. Gautier et al., (2013), for instance, found that under certain circumstances allele dropout leads to incorrect estimates of genetic diversity. Arnold et al. (2013) demonstrated through both simulations and real data that estimates of summary statistics commonly used to infer diversity and past demography from RAD data are severely affected by missing haplotypes and may show strong deviations from true values. Cariou, Duret, & Charlat (2016), finally, illustrated that allele dropout can lead to underestimation of diversity, especially in highly polymorphic species.

In light of these potentially common issues, our aims here were to develop strategies (1) to estimate genotyping error rates in RAD data, and (2) to properly incorporate the resulting genotyping uncertainty in downstream analyses to mitigate the consequences of errors. For this we present methods to estimate genotyping errors in three different ways: First, by taking advantage of available genotyping data based on a different, more reliable method (e.g. using a chip or high-depth sequencing). Second, by using independent RAD-seq replicates of individuals. And third, by assuming Hardy-Weinberg proportions among population samples. Using simulations we show that all these methods are powerful in inferring error rates even if limited samples are available. As a case study, we then applied these methods to RAD-seq data of the two widespread, genetically and ecologically divergent tree species *Populus alba* (White poplar) and *P. tremula* (European aspen) and inferred high genotyping error rates of multiple percent. By properly accounting for genotyping uncertainty, however, we obtain biologically meaningful estimates of genome-wide and local ancestry.

## Materials and Methods

### Estimation of genotyping error rates

Let us denote by *g*_*il*_ the observed genotype of individual *i* = 1, … *I*, at locus *l* = 1, …, *L*, where *g*_*il*_ = 0,1,2 reflects the number of copies of the alternative allele at a bi-allelic locus. Given per-allele genotyping error rates *ε*_*0*_ and *ε*_*1*_ at homozygous and heterozygous sites, respectively, the probabilities *P*(*g*_*il*_|*γ, ε*_0_, *ε*_1_) of observing genotype *g*_*il*_ at a locus with true genotype *γ* = 0,1,2 are given in Table 1. We next describe three strategies to estimate the genotyping error rates *ε*_*0*_ and ε_*1*_ from called genotypic data. While we present these methods in the context of Restriction site Associated DNA sequencing (RAD-seq), we stress that they are applicable to genotypic data obtained from any data source.

**Table 1.** Per-allele genotyping error model. Shown are the probabilities of observing a RAD-seq genotype given the true genotype and the per-allele genotyping error rate ε.

| True genotype | Observed genotype |  |  |
| --- | --- | --- | --- |
|  | 0 | 1 | 2 |
| 0 | $(1-\epsilon_0)^2$ | $2\epsilon_0 * (1-\epsilon_0)$ | $\epsilon_0^2$ |
| 1 | $\epsilon_1 * (1-\epsilon_1)$ | $(1-\epsilon_1)^2 + \epsilon_1^2$ | $\epsilon_1 * (1-\epsilon_1)$ |
| 2 | $\epsilon_0^2$ | $2\epsilon_0 * (1-\epsilon_0)$ | $(1-\epsilon_0)^2$ |

#### From a truth set

Consider accurate genotypes *γ*_*il*_ obtained independently for some individuals and loci. Assuming all *γ*_*il*_ to be correct and genotyping errors to be independent between sites and individuals, the likelihood of the observed genotypes ***g*** = {*g*_11_, …, *g*_*I*1_, … *g*_*IL*_} is then given by

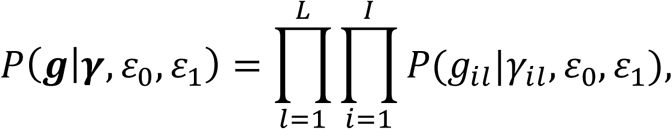

where ***γ*** = {*γ*_11_, …, *γ*_*I*1_, … *γ*_*IL*_} and *P*(*g*_*il*_|*γ*_*il*_ *ε*_0_, *ε*_1_) is given in Table 1. We obtain maximum likelihood estimates 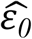 and 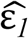 through numerical maximization (see Supporting Information).

#### From individual replicates

Consider individuals for which multiple independent sequencing experiments were conducted. Let us denote by 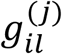 the inferred genotype of individual *i* at locus *l* in replicate *j* = 1, …, *r*_1_. The likelihood of the full data is then given by

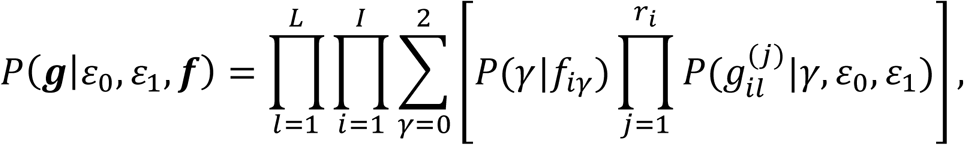

where ***g*** denotes the vector of all observed genotypes, *γ* the unobserved true genotype, *P*(*γ*|*f* _*iγ*_) = *f*_*iγ*_ the frequency of genotype *γ* among all loci of individual *i*, ***f*** = {*f*_10_, …, *f*_*I*0_, … *f*_*I*2_} and 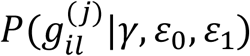 is given in Table 1. We obtain maximum likelihood estimates 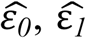 and 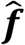 with an Expectation-Maximization (EM) algorithm as detailed in the Supporting Information.

#### From population samples

Consider individuals sampled from a random mating population such that the distribution of the true genotypes is well described by Hardy-Weinberg proportions. While the locus-specific allele frequencies *f*_*l*_ are unknown, let us assume that they follow a Beta distribution with parameters *α, β* such that *f*_*l*_∼*Beta*(*α, β*) as is expected under neutrality (Wright, 1931). The likelihood of the full data is then given by

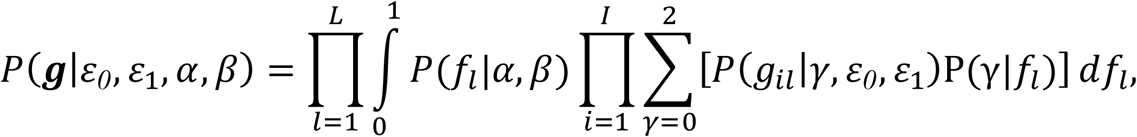

where the sum runs over the unknown true genotype *γ, P*(*γ*|*f*_*iγ*_) are the Hardy-Weinberg proportions given allele frequencies *f*_*iγ*_ and *P*(*g*_*il*_|*γ, ε*_0_, *ε*_1_), is given in Table 1. In contrast to the other two models, the computationally efficient maximum likelihood scheme resulted in biased estimates when working with population samples. We therefore resort to an MCMC approach under a Bayesian scheme (see Supporting Information) with exponential priors *ε*_0_, *ε*_1_∼Exp(*λ*) truncated at 0.5 and normal priors log(*α*), log(*β*) ∼N(*μ*, σ^2^). We used *λ =* 5, *μ* = log(0.5) and *σ*^2^ = 0.25 throughout. We also extend the model to consider data from multiple populations jointly, given that multiple samples are available for each population (see Supporting Information).

#### Set specific error rates

All methods introduced above are readily extended to jointly infer error rates for multiple sets of genotype calls that differ in their error rates, such as genotypes obtained with different sequencing depth or from different batches, for example different sequencing libraries (see Supporting Information). Inferring the error rates of all sets jointly is beneficial in the case of individual replicates or population samples, as information about hierarchical parameters such as individual genotype frequencies *f*_*i*_ = {*f*_*i*0_, *f*_*i*1_, *f*_*i*2_} or the parameters *α, β* of the Beta distribution are shared across them.

#### Recalibrating genotype likelihoods

We recalibrate genotype likelihoods by determining the likelihoods *P*(*g*_*il*_|*γ, ε*_0_, *ε*_1_) of the called genotype *g*_*il*_ for all *γ* = 0,1,2 according to Table 1 and using parameter estimates *ε*_*0*_ and ε_*1*_ obtained for the relevant set. If a truth set is available, we also calculate the empirical likelihoods *P*(*g*_*il*_|*γ*_*il*_ = *g*) across all genotypes *γ*_*il*_ = *g* of a particular set.

#### Implementation

We implemented all algorithms developed here in the open-source C++ program *Tiger* (Tools to Incorporate Genotyping ERrors), available through the git repository at https://bitbucket.org/wegmannlab/tiger. Tiger works directly off VCF files in format 4.2, which contains genotype calls encoded by the mandatory GT field. Optionally, the depth encoded by the field DP is used to bin genotype calls into different sets, which may further be stratified on a per sample basis. All methods readily handle missing data.

#### Simulations

We used simulations to assess the power of the methods introduced above to infer genotyping error rates. All simulations were generated directly under the assumed models using routines we implemented in *Tiger*. Under the truth set or replicate model, each simulated data set consisted of 50% heterozygous genotypes. Under the population samples model we drew true allele frequencies from a Beta distribution with parameters *α* = *β* = 0.7, implying about 29% of heterozygous genotypes.

To assess the bias in inferring error rates from a truth set obtained with shotgun sequencing data, we simulated sequencing data for 10^4^ truly heterozygous, 5 · 10^3^ truly homozygous reference and 5 · 10^3^ truly homozygous alternative loci at different mean depth *μ*. Specifically, we drew locus specific depths *d*_*l*_∼Pois(*μ*) from a Poisson distribution with mean *μ* and then simulated the number of observed reference alleles *r*_*l*_∼Bin(*d*_*l*_, *p*_*l*_), from a binomial distribution with probability *p*_*l*_ = 1 − *e*, 0.5, *e* for homozygous reference, heterozygous and homozygous alternative loci, respectively, assuming a sequencing error rate *e* = 0.01. We next determined the maximum likelihood genotypes 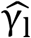 under the same model and used those to infer error rates 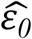 and 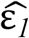 from genotypes *g*_*l*_ we simulated according to Table 1 with *ε*_*0*_ = *ε*_1_ = 0.01 or 0.1. We then generated 10^6^ replicates for each mean depth *μ* = 1, 2, …, 30 and quantified the biases 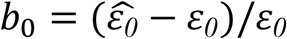 and 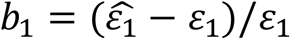.

### Application to *Populus* species

#### Study system and plant material

We generated RAD-seq data of 139 individuals of the two widespread tree species *Populus alba* and *P. tremula*, and their hybrids (*P. x canescens*) in two sets ideally suited to test the inference of error rates from a truth set and individual replicates, respectively. The first set consisted of 136 individuals (Supplementary Table S1) grown with minimal interference in a common garden established at the University of Fribourg (Switzerland) and previously genotyped with a genotyping-by-sequencing (GBS) protocol (Lindtke, Gompert, Lexer, & Buerkle, 2014). All these individuals grew from seeds collected from 15 mother trees in a natural hybrid zone in the Parco Lombardo della Valle del Ticino in Northern Italy where individuals of the two species and their hybrids grow side by side (Christe et al., 2016; Lindtke et al., 2012).

The second set consisted of four individuals for which we generated multiple replicates: a hybrid individual (F039_05) also included among the samples of the first set, a second hybrid individual (I373_A) also from the Ticino hybrid zone but grown in a common garden in Salerno, Italy, a pure *P. alba* individual (J1) from the Jalón river in the Ebro watershed (Northeast of the Iberian Peninsula), an assumed F1-hybrid tree (BET) from a population in the Tajo river headwaters (Central Iberian Peninsula). The two Iberian individuals were previously genotyped using microsatellites (Macaya-Sanz et al., 2011).

#### DNA extraction and RAD sequencing

For all samples, DNA was extracted from 15-20 mg of silica-dried leaf material with the Qiagen DNeasy Plant Mini Kit (Valencia, CA). The concentration of DNA was measured with a Qubit 2.0 Fluorometer using the dsDNA HS assay kit (Invitrogen), and its integrity verified with electrophoresis on 1.5% agarose gels (1X TBE). Concentrations were standardized to 20 ng/μl and individual samples were submitted for RAD library preparation and sequencing to Floragenex (Eugene, OR). There, all extractions of the individuals of the first set, as well as the replicate extractions of F039_05 and I373_A, were processed (together with additional samples prepared in the same way), in five libraries of 95 individuals each. These libraries were prepared according to Floragenex’ standard commercial protocol: genomic DNA was digested with the restriction endonuclease *Pst*I (Christe et al., 2016; chosen according to previous studies on these species - Stölting et al., 2013) and RAD libraries were prepared with a method similar to the one described in Baird et al. (2008). This protocol included 18 PCR cycles, after which DNA fragments ranging from 300 to 500 bp were retained. All five libraries were sequenced in a single run on an Illumina HiSeq 2500 instrument, but on individual lanes. This data is available in the Sequence Read Archive through bioproject PRJNA528699.

Following the same protocol, an additional library was generated and sequenced by Floragenex in a separate experiment, consisting of two and three replicate extractions of J1 and BET, respectively, as well as extractions from offspring of a controlled cross between them. This data is available in the Sequence Read Archive through bioproject PRJNA528706.

#### Bioinformatic data processing

We assigned reads to individuals or replicates with *fastq-multx* (eautils; Aronesty, 2011), allowing one mismatch in the 15 bp including barcode and restriction site. Read quality was checked with *FastQC 0.10.1* (S. Andrews, 2010) and low quality bases and reads were removed with *condetri* v.2.3 (Smeds & Künstner, 2011) using default parameters, except for the options *-hq* (high quality threshold) and *-lfrac* (maximum acceptable fraction of bases after quality trimming with quality scores lower than the threshold *-lq*), for which a value of 15 and 0.1 were chosen, respectively.

High-quality reads were aligned against the *P. tremula* mitochondrial reference sequence (Kersten et al., 2016) and against the nuclear reference genome of *P. trichocarpa* (Ptrichocarpa_210_v3.0; Tuskan et al., 2006) using *Bowtie2 2.3.0* (Langmead & Salzberg, 2012) with “end-to-end” and “very sensitive” settings. Reads with mapping quality lower than 20 were discarded using *samtools 1.3* (Li et al., 2009) and read group information was added with *picard tools 1.139* (http://broadinstitute.github.io/picard). We then used the tools *TargetCreator* and *IndelRealigner* of *GATK 3.8* (DePristo et al., 2011) to realign around indels, and recalibrated base quality scores for each individual using the method by Kousathanas et al. (2017) implemented in *ATLAS* (Link et al., 2017) on mitochondrial sequences. This method does not require *a priori* information on genotypes and instead learns base qualities from haploid regions while integrating over genotype uncertainty. Finally, we called genotypes with *UnifiedGenotyper* in *GATK 3.8* (DePristo et al., 2011).

To then only retain reliable sites for comparison, we filtered resulting variants using *vcftools* (Danecek et al., 2011) and custom R scripts: first, we removed sites with an average depth across individuals ≥24 (the 98.7% quantile) to exclude potentially paralogous loci. Second, we only kept Single Nucleotide Variants (SNVs) with at most two segregating alleles. Third, we removed all variant sites within 5 bp of an indel to avoid SNVs due to misalignments.

#### Truth Set

Genotypes of the 136 individuals grown in the common gardens were previously obtained (Lindtke et al., 2014) with a genotyping-by-sequencing (GBS) protocol very similar to the ddRAD protocol (Peterson et al., 2012) and using the restriction enzymes *EcoR*I and *Mse*I. Importantly, (Lindtke et al., 2014) generated sequencing data also for the 15 mother trees and used sibships in a Bayesian approach to infer genotypes while accounting for familial relationships.

To compare these high quality genotypes to those obtained from our own RAD-seq experiment, loci covered in both studies had to be identified first. Since Lindtke et al. (2014) used an older *P. tremula* reference, we extracted from this reference windows of 201 bp around each locus in the GBS data set (100 bp on either side). We then mapped these extracted sequences against the *P. trichocarpa* reference with *Bowtie2 2.3.0* (Langmead & Salzberg, 2012) with “end-to-end” and “very sensitive” settings and retained only those sequences that mapped uniquely with quality of 20 or more. We then kept all loci overlapping between the two data sets, but removed four loci for which different alternative alleles were called. To ensure high accuracy of the GBS data, we restricted all comparisons to genotypes called with a posterior probability ≥99% by Lindtke et al. (2014).

#### Estimating genotyping error rates

We estimated genotyping error rates from our RAD-seq data from both individual replicates as well as through the comparison to the truth data described above. Since our data does not reflect random samples of a population, we could not benefit from the population sample method introduced above.

#### Quantifying allele dropout

We obtained a rough estimate of allele dropout by remapping the raw GBS data against the *P. trichocarpa* reference with *bwa-mem* (Li, 2013), identifying the most likely alternative allele and calling genotypes using the task majorMinor of *ATLAS* (Link et al., 2017) for all sites within *Pst*I cut sites, which we identified in the *P. trichocarpa* reference using REDseq (Chen *et al.*, 2014). We then quantified the fraction of cut sites across individuals with at least one heterozygous genotype, out of all cut sites that were covered at depth ≥ 3x at all sites.

#### Estimating of genome-wide and interspecific ancestry

For each of the 136 common garden samples we estimated the genome-wide proportion of *P. alba* ancestry (genome-wide ancestry *q*) and the proportion of the genome heterozygous for the two ancestries (interspecific ancestry *Q*_*12*_) using *entropy* (Gompert et al., 2014), a program that implements a model similar to the admixture model in *structure* (Pritchard, Stephens, & Donnelly, 2000). In contrast to *structure*, however, *entropy* can also make use of uncertain genotypes from low depth sequence data by working directly with genotype likelihoods, rather than genotype calls. Here we ran *entropy* on the raw genotype likelihoods provided by UnifiedGenotyper, as well as on genotype likelihoods recalibrated using empirical likelihoods estimated from the comparison to a truth set, excluding sites with >50% missing data in both cases. To stratify the estimates and have sufficient observations to estimate these probabilities reliably, we considered five RAD-seq depth classes: 1-3, 4-7, 8-15, 16-31 and ≥32.

#### Inference of locus-specific ancestry

To infer locus-specific ancestry, we ran *RASPberry* (Wegmann et al., 2011), which implements a Hidden Markov Model (HMM) to explain haplotypes of admixed individuals as a mosaic of provided reference haplotypes for each species. We obtained suitable reference haplotypes by phasing previously characterized pure *P. alba* and pure *P. tremula* individuals (51 each) from the Italian, Austrian and Hungarian hybrid zones (Christe et al., 2016) using *FastPhase* (Scheet & Stephens, 2006), building input files with *fcGENE* (Roshyara & Scholz, 2014). For use in *RASPberry*, individuals in the reference panels were not allowed to have missing data. We thus restricted the comparison to only the SNVs covered in all parental individuals.

To compute HMM transition probabilities in *RASPberry*, we used a default recombination rate of 5 cM/Mb as estimated by Tuskan et al. (2006) in *P. trichocarpa* and the estimates of the genome-wide ancestry *q* for each sample obtained with *entropy* from the error corrected data. For most other parameters we used previous estimates for *P. alba* and *P. tremula* hybrid zones (Christe et al., 2016), but scaled these as proposed by Wegmann et al. (2011) to reflect the size of the reference panel. These include the ancestral population recombination rates (315 and 900 for *P. alba* and *P. tremula*, respectively), mutation rates (0.00185 and 0.00349, respectively), the miscopying rate (0.01) and the miscopy mutation rate (0.01). However, we set the time since admixture to five (rather than one) to reflect the different sampling strategy.

To account for genotyping errors, we determined a per-allele genotyping error *ε* as the weighted average across depth-specific error rates obtained under the truth set model, with the constraint *ε*_0_ = *ε*_1_ and weights reflecting the distribution of depth in the sites included in the analysis. We then added this error rate estimate to the population specific and miscopy mutation rates. Under the *RASPberry* copying model, these parameters control the rate at which the sample genotypes differ from the reference haplotype from which the sample is copying. That rate thus depends on the reference panel size, but also on genotyping errors.

We called ancestry segments as any stretch on a chromosome within which the posterior probabilities for a particular ancestry (homozygous *P. alba*, heterozygous ancestry or homozygous *P. tremula*) was >0.5 at all SNVs, and measured its length from the first to the last SNV.

## Results

### Power to infer genotyping error rates

Simulations suggest that a few thousand loci are sufficient to accurately estimate genotyping error rates even from just a few samples (Figure 1). Due to the extra information provided, the smallest estimation errors were obtained when using a truth set: >90% of all estimates fell within a range from half to two-fold the true value (Q2, e.g., within [0.005, 0.02] for a true error rate of 0.01) if estimated genotypes were compared at 200 and 400 sites for _*0*_ and ε_*1*_, respectively. Similar accuracy was achieved under the population samples model as soon as 5,000 sites were used.

**Fig. 1:**
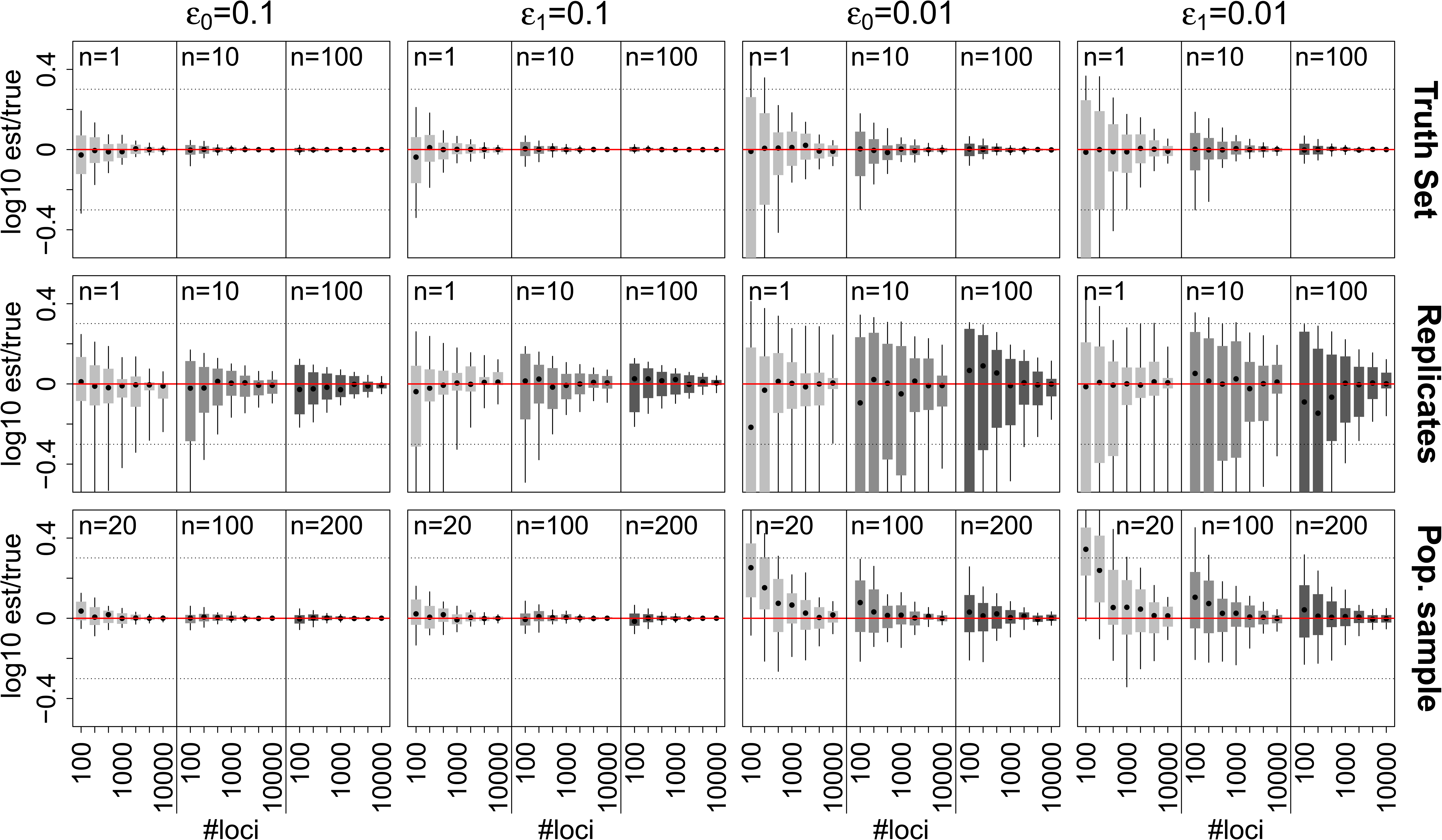
Accuracy of error rate estimates. Shown are the estimated error rates relative to the true error rates (red line) of 100 replicates for different samples sizes n (shown on top), the two error rates 0.1 and 0.01 (first and last two columns, respectively) and different numbers of unlinked loci. The horizontal dashed lines indicate Q2, the interval within which an estimate is less than a factor of two away from the true value (i.e. within half and two times the true value). Simulations were generated for the truth set (top row), the individual replicate (two replicates per sample, middle row) and population samples models (bottom row).

The lowest accuracy was observed under the replicates model, especially if error rates were low. Using 5,000 sites, for instance, all estimates were within Q2 for a true value of *ε*_0_ = *ε*_1_ = 0.1, but only slightly above 80% of all estimates for a true value of *ε*_0_ = *ε*_1_ = 0.01. This is readily explained by the fact that only very limited information about the true genotype is available: if the two replicates differ in their genotype, it is not clear which one is correct. Consequently, the accuracy of inference is much increased if more than two replicates are available per individual (Supplementary Figure S1). Nonetheless, even small error rates can be estimated relatively accurately, as >90% of all estimates for true value of *ε*_0_ = *ε*_1_ = 0.01 fell within Q2 as soon as 10^4^ or more comparisons were used.

Importantly, the accuracy is a function of the fraction of truly heterozygous genotypes in the data set, with the accuracy of ε_*1*_ increasing with more heterozygous calls, and that of *ε*_*0*_ and ε_0_ decreasing. Here we simulated 50% heterozygous genotypes for the truth set and replicate models and about 29% heterozygous genotypes for the population sample model. If the number of heterozygous genotypes were lower, proportionally more sites would be needed to achieve the accuracy presented here. If 20% of all genotypes were heterozygous, for instance, around 10^5^ sites are required under the replicate model with two replicates per sample.

### Using Whole-Genome Sequencing to Establish a Truth Set

As shown above, genotyping error rates are inferred particularly accurately when comparing calls to a truth set. One possibility to establish such a truth set is to shot-gun sequence a subset of individuals and to infer genotypes from this data. This is also feasible in case no reference genome is available as the shot-gun reads can be mapped directly against the RAD-loci. But since genotyping errors in the truth set results in an overestimation of genotyping error rates, sufficiently high depth must be used. Here we used simulations to assess this bias for the case of shot-gun sequencing one individual at 10^4^ homozygous and 10^4^ heterozygous RAD loci.

As shown in Figure 2 for *ε*_0_ = *ε*_1_ = 0.01, genotyping error rates may be overestimated considerably unless the depth used to establish truth set was relatively high. At a mean depth of 10x, for instance, the median relative bias was 1.92 and 0.50 for *ε*_0_ and *ε*_1_, respectively, and hence on the same order as the inferred error rate. At a mean depth of 30x, the median overestimation dropped below 0.01 in both cases. Importantly, larger quantiles drop much slower, suggesting that a minimal bias is hard to overcome even with very high depths. Noteworthy, we observed a stronger bias for *ε*_0_ than *ε*_1_ at all depths, a result of more often calling truly heterozygous sites homozygous than vice versa from next-generation sequencing data. Results were virtually identical for the case of *ε*_0_ = *ε*_1_ = 0.1, but scaled due to the larger true error (Supporting Figure S2).

**Fig. 2:**
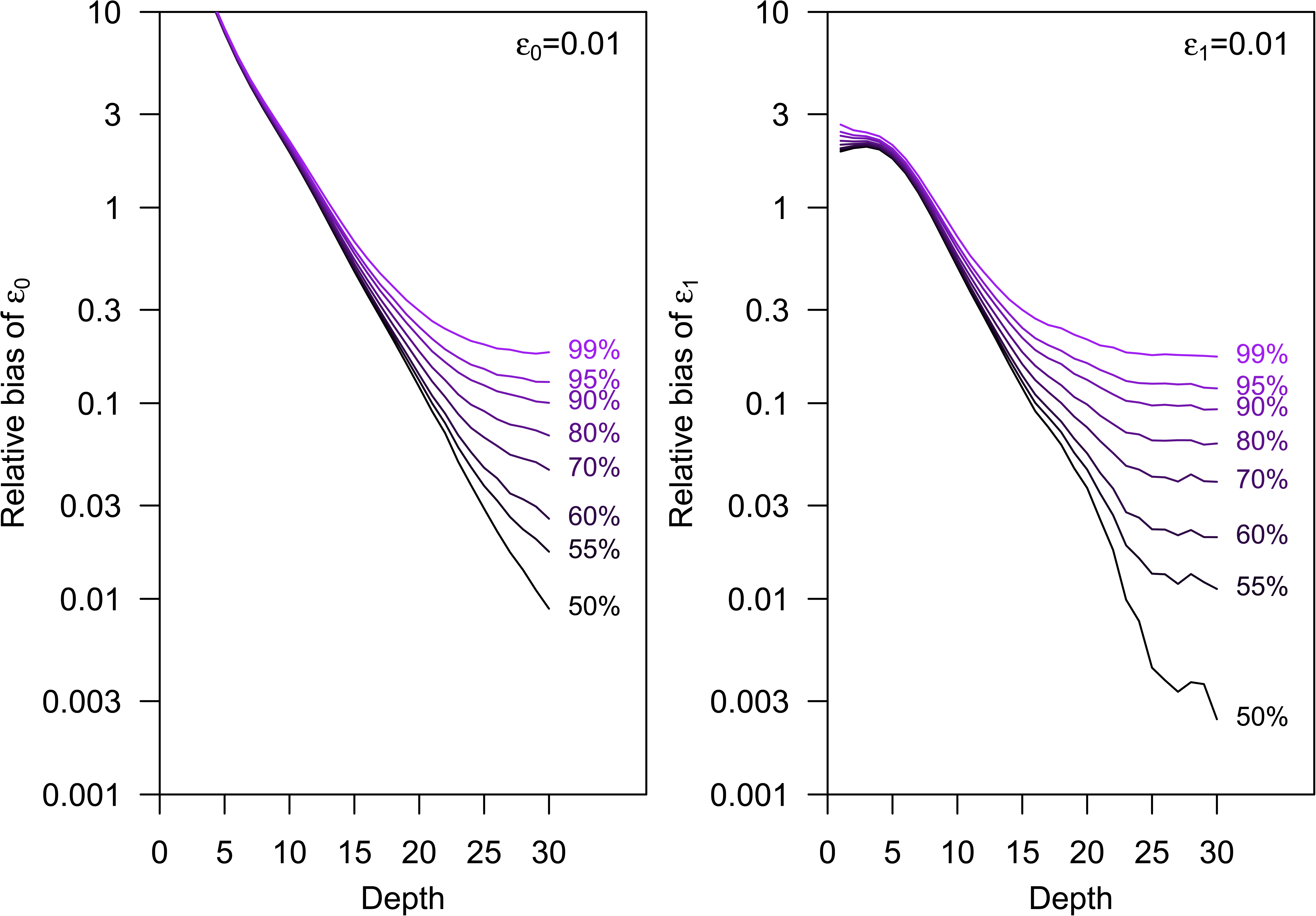
Distribution of the relative bias when inferring error rates from a truth set generated with next-generation sequencing data with a specific mean depth. Each line indicates a quantile m such that a smaller bias was observed in a fraction m of 10^6^ simulations of 10^4^ homozygous and 10^4^ heterozygous loci with *ε*_0_ = *ε*_1_ = 0.01.

### High genotyping error rates in RAD-seq

We next used our inference methods to quantify genotyping errors from our RAD-seq experiment of 137 individuals of the two widespread tree species *Populus alba* and *P. tremula*, and their hybrids (*P. x canescens).* On average, our experiment resulted in 831,161 (sd 153,434) reads per sample that passed quality trimming and mapped against the reference genome of *P. trichocarpa* with mapping quality ≥20. From those, we called 529,305 Single Nucleotide Variants (SNVs), after removing multi-allelic sites, those with excessive depth, indels and variant sites around indels. We estimated per-allele genotyping error rates from these SNVs through a comparison with previously published, high-quality genotypes (truth set), and from multiple replicate libraries sequenced for a subset of our samples (replicates).

#### Truth set

We estimated per-allele genotyping errors by comparing genotype calls from our RAD-seq experiment to those of a previously published GBS dataset (Lindtke et al., 2014) for 136 individuals present in both studies. In total, 7,426 SNVs overlapped between experiments, at which we could use a total of 16,610 genotype comparisons. Of those, only 69.9% matched, with matching rates increasing with RAD-seq depth (Figure 3A). Strikingly, RAD-seq genotypes were much less often heterozygous than GBS genotypes (Figure 3B), especially at low depth. In line with these observations, we inferred per-allele genotyping error rates >10% whenever RAD-seq depth was ≤ 35 and when using a model assuming a single error rate (*ε*_*0*_ = *ε*_1_), driven by an exceptionally high error rate at truly heterozygous sites *ε*_1_ (Figure 2C).

**Fig. 3:**
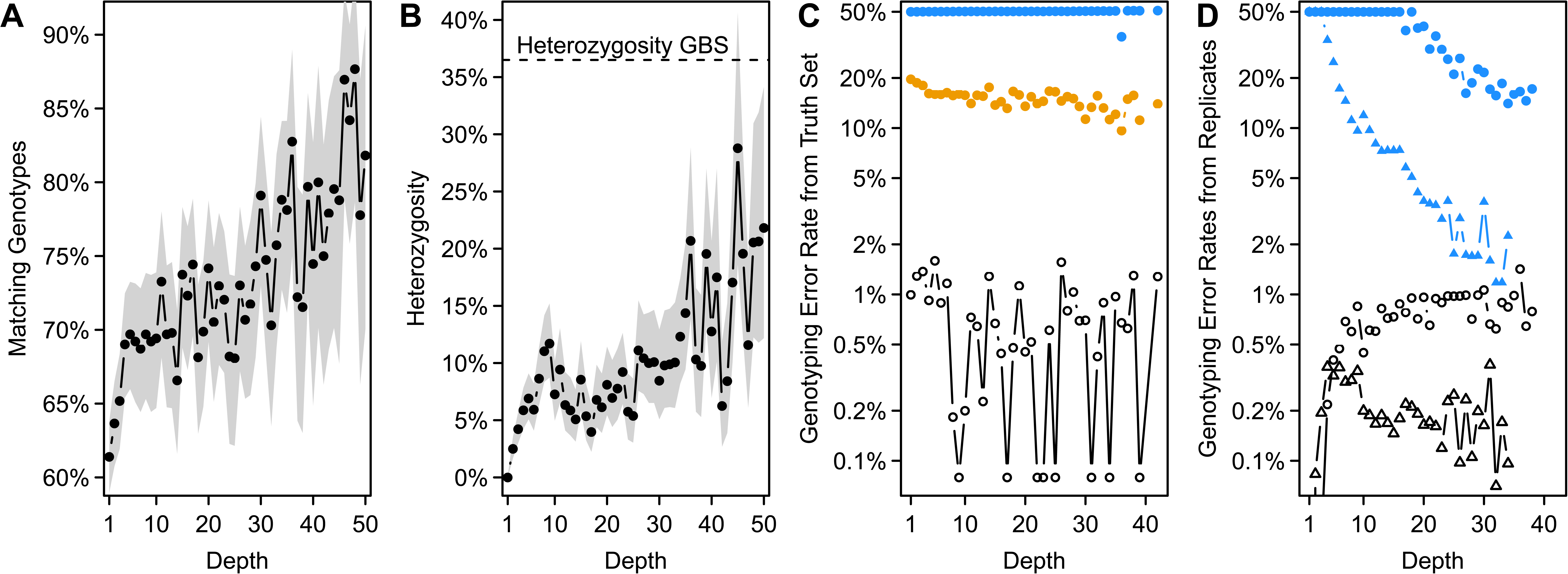
Estimates of genotyping errors in RAD-seq data. **A**. Maximum likelihood estimate (MLE) of the frequency of genotypes estimated from GBS and RAD-seq data that match as a function of RAD-seq sequencing depth. The gray region indicates the range of frequencies within two log-likelihood units of the MLE. **B.** MLE estimate of the frequency of heterozygous calls with RAD-seq among all calls of a particular depth overlapping with the GBS data. Gray region as in A. The heterozygosity observed among all overlapping GBS genotypes is given as dashed line. **C**. Per-allele RAD-seq genotyping error rates for homozygous (*ε*_0_, black open symbols) and heterozygous (*ε*_1_, blue filled symbols) genotypes estimated using a GBS truth set, limited to depth class with at least 100 comparisons. Estimates for a model assuming ε_0_ = *ε*_1_ are shown with yellow closed circles. **D.** Per-allele RAD-seq genotyping error rates obtained from two replicates of F039_05 and I373_A each (circles) and two and three replicates from J1 and Bet, respectively (triangles), limited to depth class with at least 1000 sites. Colors as in C.

These are surprisingly high error rates, particularly when considering that sequencing error rates of Illumina machines are estimated at <1% (Nielsen, Paul, Albrechtsen, & Song, 2011). Importantly, the bias towards homozygous genotype calls is not simply explained by low depth. Indeed, RAD-seq still resulted in less than half as many heterozygous calls at depths ≥40x, which are usually considered more than sufficient for accurate genotype calling (Nielsen et al., 2011). Instead, our results suggest an inherent bias in the RAD-seq data analyzed here.

We confirmed our suspicion by analyzing allelic imbalance at sites thought to be heterozygous based on the GBS genotypes, limiting the analyses to genotypes covered at up to 50x to avoid spurious alignments (Supplementary Figure S3). Only a single allele was for instance observed at 69% of all heterozygous genotypes with depth ≥ 5x, a fraction that dropped to 60% and 52% for genotypes with depth ≥ 20x and depth ≥ 30x, respectively, indicative of allelic dropout at a large scale.

However, our estimates rely on the assumption that the GBS data reflect true genotypes. This is based on good evidence: First, Lindtke et al. (2014) additionally sequenced the mother trees of all individuals considered here and estimated posterior genotypes using a hierarchical ancestry model that incorporated familial relationships with mothers and among siblings. These updated estimates correlated with the raw maximum likelihood genotype estimates ignoring familial data at 0.985 and differed from those in <0.02% of all calls. Second, we restricted this comparison to GBS genotypes with a posterior probability ≥99%. Third, the fraction of concordant genotype calls between the GBS and RAD-seq data increased with RAD-seq depth. If the mismatches were driven by errors in the GBS data, no such dependence should be observed.

#### Replicates

We next benefitted from two sets of replicate libraries to estimate per-allele genotyping error rates. The first set consisted of two replicate libraries of each of two individuals (F039_05 and I373_A) sequenced along all other samples. Error rates estimated from that data corroborated the conclusion obtained from the comparison with GBS data (Figure 3D): error rates at truly homozygous sites (*ε*_0_) were on the order of 1% or less, and those at truly heterozygous sites (*ε*_1_) were equal or close to 50%, which is the largest value possible under our model.

In contrast to the estimates obtained in comparison to GBS genotypes, the error rates at truly heterozygous sites (ε) dropped to about 20% at high depth (≥20x). This difference might in part be driven by errors in the GBS data slightly inflating error rate estimates. However, given the high quality of the GBS data, it appears more likely that the error rates from replicates are underestimated. Polymorphisms in restriction cut sites or unequal PCR amplification rates of alleles, for instance, affect replicates systematically, while the statistical inference must assume independence of errors between replicates.

To verify that high error rates are not specific to the RAD-seq run performed on these hybrids, we carried out a second RAD-seq experiment including two and three replicates of a pure *P. alba* individual (J1) and a putative F1 *P. alba* x *P. tremula* hybrid (BET), respectively. This experiment resulted in 694,031 (sd 187,957) reads per sample that passed quality trimming and mapped against the reference genome of *P. trichocarpa* with quality ≥20. Per-allele error rates estimated from this data were indeed lower, with error rates at truly homozygous sites (ε_0_) on the order of 0.2% and those at truly heterozygous sites (ε_1_) starting out at 50% and dropping to about 3% at a depth of 25x (Figure 3D). However, these lower errors still point to a particular issue in calling heterozygous genotypes: of all truly heterozygous sites with depth ≥25x in our data, >5% are expected to be called homozygous. (The lower output of this sequencing experiment does not allow us to make reliable statements for higher depths).

### Estimation of genome-wide and interspecific ancestry

We investigated the impact of genotyping errors in our RAD-seq data on the inference of genome-wide ancestry *q* as well as interspecific ancestry *Q*_*12*_, which reflects the proportion of loci of heterospecific ancestry. We estimated these ancestry components with *entropy* from 230,805 SNVs, and compared them to estimates from GBS obtained by Lindtke *et al.* (2014).

The model implemented in *entropy* accounts for the genotyping uncertainty reflected in the genotype likelihoods. However, the raw genotype likelihoods obtained from our RAD-seq data are misleading: for sites with considerable depth, the RAD-seq genotype likelihoods often suggest almost certainty for wrong genotypes. Of all genotypes wrongly called as homozygous at depth ≥30x (judged by the comparison to GBS data), 90% had a variant quality of 77 or more. (A variant quality of 77 implies that it is more than 5 billion times less likely to observe the obtained data from a heterozygous than homozygous site).

As a result, ancestry estimates differed considerably between the GBS and RAD-seq data sets (Figure 4). Interestingly, the estimates of the genome-wide ancestry *q* were much less affected by genotyping errors than the estimates of the interspecific ancestry *Q*_*12*_. This is readily explained, however, by the directionality of the most common error, which is to wrongly infer homozygous genotypes at heterozygous sites. We found these errors do result more frequently in a homozygous reference than homozygous alternative call (65.0%), particularly at low depth (84.5% at depth ≤5x, 49.9% at depth ≥20x). But this did not introduce a bias in *q* towards one of the species since we used the sequence of the outgroup *P. trichocarpa* as reference. The estimates of *Q*_*12*_, however, were very sensitive to an underestimation of heterozygosity.

**Fig. 4:**
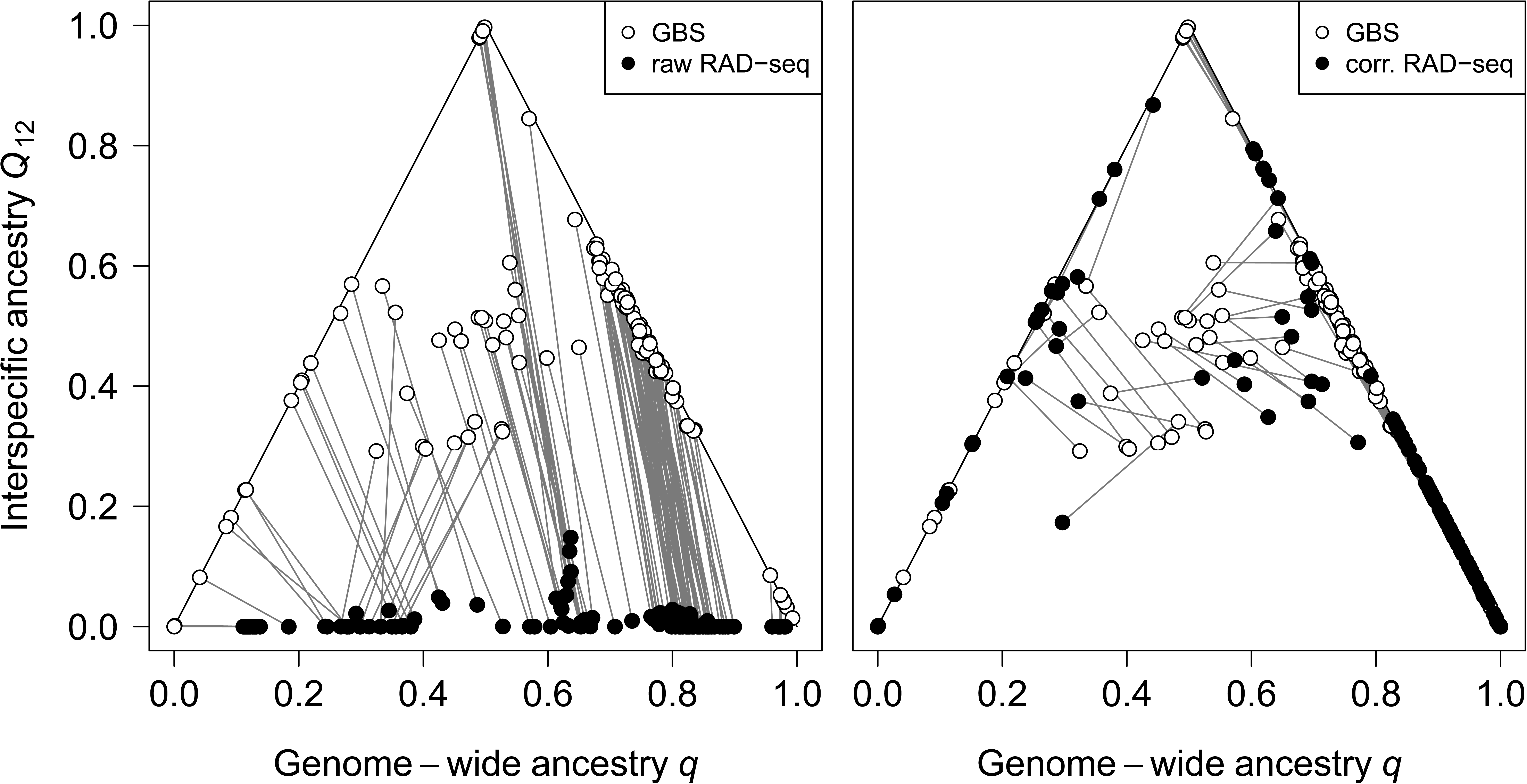
Comparison of genome-wide (*q*) and interspecific (Q_12_) ancestry estimates of 136 individuals obtained from GBS data (Lindtke et al., 2014) and estimates from RAD-seq data (black circles) using either the raw (left panel) or corrected (right panel) genotype likelihoods. Grey lines connect values obtained for the same individual.

To improve these estimates, we propose to directly account for the elevated genotyping error rates in RAD-seq data by adjusting the genotype likelihoods according to the observed genotyping uncertainty. By treating the genotype calls as data, we can determine the probabilities *P*(*g*|*γ*) of observing a RAD-seq genotype call *g* given the true genotype *γ* either by using estimates of the per-allele error rates *ε*_0_, *ε*_1_, or empirically from the comparison to a truth set. Using the latter approach on our data (individually for each depth) resulted in estimates of *q* and *Q*_*12*_ that were much closer to those obtained by Lindtke *et al.* (2014; Figure 4).

Some differences between the point estimates of *Q*_*12*_ remain, likely due to 1) differences in the models and their information-sharing among individuals, as Lindtke *et al.* (2014) used information about maternal plants and sibships as part of the model, and 2) the extent to which information in uncertain genotypes was outweighed by hierarchical prior probabilities related to ancestry. Nonetheless, these results demonstrate the importance of accounting for uncertainty in genotyping data since the estimates of interspecific ancestry with and without correction lead to a very different biological interpretation: With correction, the hybrid individuals appear to be mostly early generation hybrids, suggesting meaningful reproductive isolation between the species. Without correction, the large number of individuals with low *Q*_*12*_ but intermediate values of *q* suggests considerable gene flow between the species, an interpretation at odds with recent work (Christe et al., 2016, 2017; Macaya-Sanz et al., 2011).

### Inference of locus-specific ancestry

We next evaluated the impact of genotyping errors on local ancestry inference. Hybrid zones between *P. alba* and *P. tremula* are dominated by pure parental individuals and early hybrids (mostly F1), with only few adult recombinant hybrids (Christe et al., 2016; Lindtke et al., 2014). We chose ten individuals among our samples spanning that spectrum according to *q* and *Q*_*12*_ values from Lindtke *et al.* (2014): a putatively pure *P. alba* individual (F039_01), a putatively pure *P. tremula* individual (F030_01), two putative backcrosses to *P. alba* (F020_04 and F032_08), two putative backcrosses to *P. tremula* (I345_02 and I345_03), two putative F1 hybrids (I373_03 and F030_05) and two putative hybrids of later generations (F022_03 and F026_05). We then used *RASPberry* (Wegmann et al., 2011) to infer local ancestry along chromosomes 1 through 5, restricting our inference to the 6,445 SNVs that did not have missing data in the parental reference haplotypes we took from Christe *et al.* (2016).

In line with these expected simple ancestry make-ups, we inferred many large ancestry blocks often spanning almost entire chromosomes (Figure 5). Surprisingly, however, we inferred most of these blocks to be of homospecific ancestry, and also inferred many short segments, which are difficult to reconcile with the putative ancestries of our samples (Figure 5). As an example, consider the individual I373_03 in Figure 5 that was classified as an F1 hybrid by Lindtke et al. (2014), but for which we inferred homospecific ancestry blocks for both parental species. Such artifacts could arise from the reference panels being too small to properly reflect the haplotypes found in our hybrid individuals, large gaps between neighboring SNVs limiting the power of the HMM implemented in *RASPberry*, but most likely by genotyping errors towards homozygous genotypes.

**Fig. 5:**
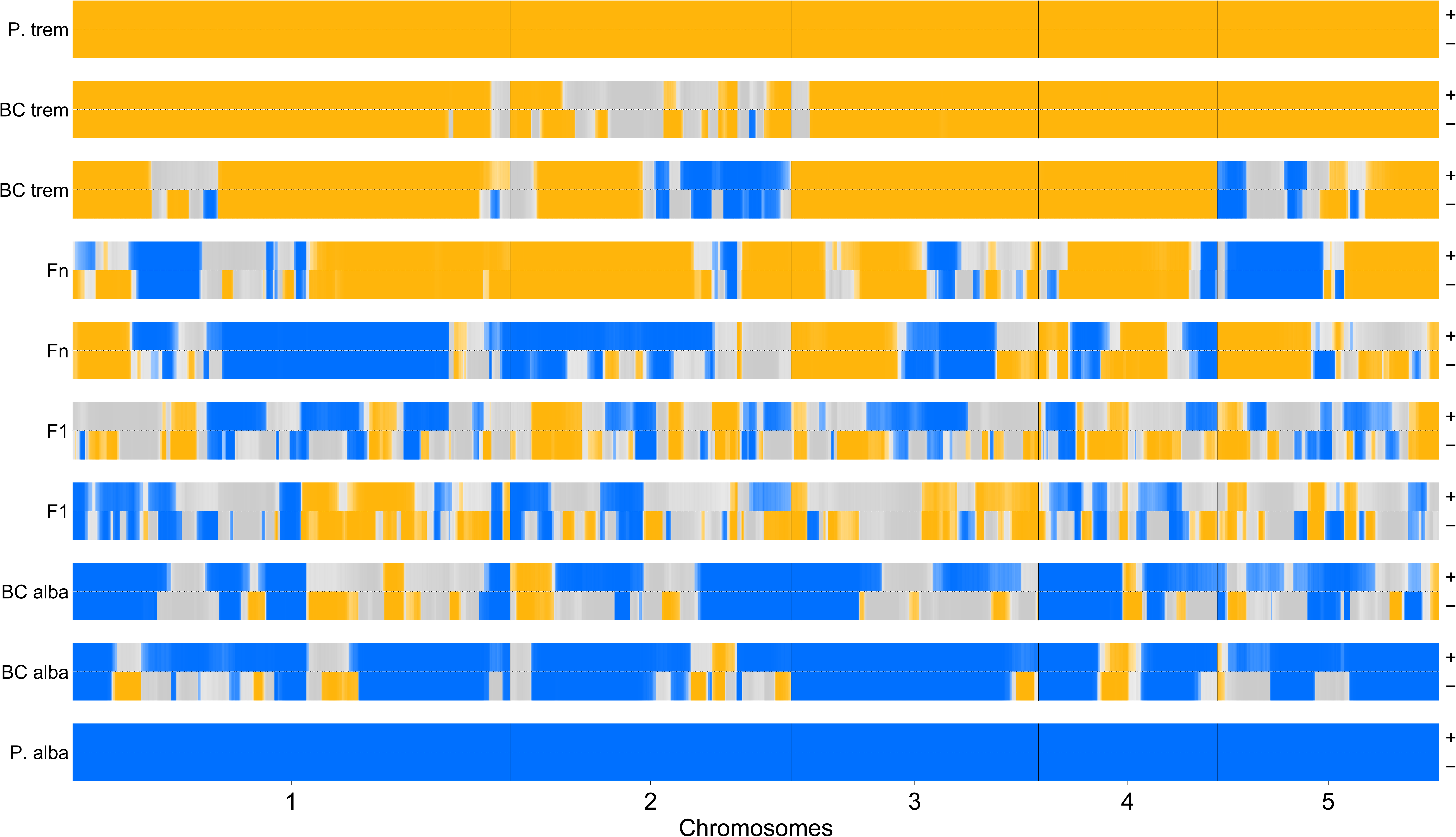
Comparison of local ancestry patterns on chromosomes 1-5 for (top to bottom) a putatively pure *P. tremula* individual (*P. trem*, F030_01), two putative backcrosses to *P. tremula* (BC trem, I345_02 and I345_03), two putative hybrids of later generation (Fn, F022_03 and F026_05), two putative F1 hybrids (F1, I373_03 and F030_05), two putative backcrosses to *P. alba* (BC alba, F020_04 and F032_08) and a putatively pure *P. alba* individual (*P. alba*, F039_01). For each individual, RASPberry results with (+) and without (-) correction are shown. Blue represents *P. alba* ancestry, orange *P. tremula* ancestry and grey heterospecific ancestry, with darker shades showing higher confidence in the ancestry estimates. To facilitate visualization, only sites with data are shown.

While *RASPberry* does not account for genotyping uncertainty via genotype likelihoods, the implemented copying-model allows for “mutations”, or differences between the observed genotype of an admixed individual and the reference haplotypes from which it is copying. We thus repeated the inference by adding an estimate of the average per-allele genotyping error rate obtained under the truth set model (13.7%) to the mutation rate parameters to account for the high genotyping error in our RAD-seq data (Figure 5). Accounting for genotyping errors indeed improved our estimates. For the putative backcrosses, for instance, we called fewer segments homozygous for the “wrong” ancestry (11 versus 34) and these covered a smaller fraction of the first five chromosomes (6.1% versus 11.1% of the parts at which ancestry could be called). Similarly, we called a higher fraction of the first five chromosomes to be of heterozygous ancestry (45.0% versus 37.2% of the parts at which ancestry could be called). While these results corroborate the importance of accounting for the true uncertainty in genotypes in downstream analysis, they also illustrate that a method accounting for a uniform error fails to fully mitigate the bias against heterozygous genotypes present in our RAD-seq data sets.

## Discussion

### High Genotyping Error Rates in RAD-seq

Here, we report high genotyping error rates in two independent RAD-seq data sets. We obtained these estimates by comparing RAD-seq calls of two independent experiments, either to published genotype calls for the same individuals (Lindtke et al., 2014) obtained with a different sequencing method (GBS), or to calls obtained from independent replicates of the same individuals. Both approaches provide evidence for high per-allele genotyping errors of several percent and show that the RAD-seq data analyzed here has a strong bias towards calling homozygous genotypes at heterozygous sites that is not overcome with higher sequencing depth.

While these results do not directly generalize to other RAD-seq studies or other RAD-seq protocols, the few estimates of genotyping errors reported to date agree with the high estimates we obtained here. Luca et al. (2011), for instance, compared genotypes of human samples obtained with a technique similar to RAD to those available in a public database, and estimated that between 6.3 and 9.7% of heterozygous sites were called as homozygous. Similarly, Mastretta-Yanes et al. (2015) found that, depending on the parameter settings chosen for the *de-novo* assembly, between 5.9 and 8.8% of alleles were not concordantly called between replicates.

Several factors could explain the high genotyping error and the lack of heterozygous genotypes frequently reported in RAD-seq data. Of major concern is the issue of allele dropout due to polymorphisms in restriction cut sites and the “loss” of one allele at heterozygous sites (Andrews et al., 2016; Davey et al., 2013; Puritz et al., 2014). We obtained a rough estimate of the potential for allele dropout in our RAD-seq data by focusing on *Pst*I cut sites in the *P. trichocarpa* reference genome that were covered by a depth of ≥ 3x with GBS data. Among those, we then identified cut sites that contained a heterozygous genotype call at at least one site. Of the 550 cut sites with sufficient data, 89 (16.2%) harbored at least one heterozygous genotype call, affecting 24 (14.8%) of the 162 loci for which at least one individual had data. While these estimates are rough, they do suggest that allele dropout may have contributed significantly to genotyping error in our RAD-seq data. Indeed, they suggest that about one third of all RAD-seq loci could have been affected by a cut site polymorphism on at least one fragment end, leading to allele drop-out in case of a linked polymorphism within the locus.

However, allele dropout is unlikely to account for all genotyping errors observed, as indicated by the many sites with strong allelic imbalance but without complete dropout (Figure S3). Alternatively, one allele might for instance not have been sequenced or sequenced only at very low depth because of differences in fragment length that can lead to amplification bias, less efficient shearing, or loss in size selection. This is a likely explanation, since Davey and colleagues (2013) showed that there is a high correlation between read depth and fragment length and the protocol used here included 18 cycles of PCR. Similarly, differential efficiency of PCR among alleles could have masked one allele (i.e. PCR duplicates), causing it to be represented in a very low number of reads (Schweyen et al., 2014). However, PCR duplicates were recently reported to be of only minor concern in RAD-seq studies, at least if only up to 12 PCR cycles were used (Euclid 2019). Finally, PCR errors, in conjunction with differential amplification, might have resulted in homozygous alternative calls at homozygous reference sites (or vice versa) we observed rarely, but even at high depth (Figure S3).

### Accounting for Genotyping Error Rates

Several bioinformatic solutions have been suggested to mitigate the apparent biases in RAD-seq. Both Arnold et al. (2013) and Gautier et al. (2013) recommend the comparison of read depth across sites, to identify loci likely exhibiting allele dropout. In our case, however, depth varied substantially across sites, because of PCR duplicates or stochastic events, rendering such an approach difficult. Davey et al. (2013) also noted that alleles present in two copies at homozygous sites have higher depth compared to alleles present in single copy at heterozygous loci, but read depths for the two sets of alleles overlap, inhibiting the accurate detection of loci with allele dropout by using depth alone. To improve upon this, Cooke et al. (2016) developed a method to infer the likelihood of observing allele dropout at a site on the basis of the sequencing depth of each sample, and suggested to ignore sites where this likelihood is high. Finally, it was also suggested to discard any locus with a missing genotype, since this might indicate a polymorphism in the restriction site. However, in many studies with moderate depth, including ours, the amount of missing data prevents the adoption of such drastic solutions. In summary, all these filtering suggestions result in a massive reduction in usable loci, and hence further accentuate the already limited genome-wide coverage of reduced library techniques such as RAD.

As a model-based alternative, we propose here to properly account for the high genotyping errors in downstream analysis. A first such attempt was proposed by Cariou et al. (2016), who developed an Approximate Bayesian Computation (ABC) method to estimate genetic diversity while accounting for allele dropout, but found this method not to be accurate under elevated levels of diversity. A more general solution, we believe, is to make use of the large number of recently developed tools that do not require genotype calls but rather work directly from genotype likelihoods to account for uncertainty in the data (Fumagalli, Vieira, Linderoth, & Nielsen, 2014; Jørsboe, Hanghøj, & Albrechtsen, 2017; Korneliussen, Albrechtsen, & Nielsen, 2014; Kousathanas et al., 2017). Using such tools minimizes the necessity to filter data stringently and is readily applied to low-depth data (Nielsen et al., 2011).

For such methods to work properly, however, the genotype likelihoods need to accurately reflect the uncertainty in genotypes. While all modern genotype callers also calculate genotype likelihoods, these do not reflect biases specific to individual sequencing protocols such as RAD-seq, as we illustrated here, and must thus be recalibrated. Here we propose three recalibration strategies: If accurate genotype calls are available for a subset of the individuals and markers, empirical genotype likelihoods can be obtained by comparing those to calls from a reduced representation sequencing experiment. Alternatively, individual replicates or population samples may be used to infer per-allele genotyping error rates, from which recalibrated genotype likelihoods are readily calculated. Tools for these strategies are available through the software *Tiger*, which also accounts for sequencing depth as an additional covariate. While we found sequencing depth to be a particularly important predictor, the model is also readily extended to additional covariates such as the raw genotype likelihood or genotype call, which might provide additional information about genotyping error rates.

However, all these strategies might be biased. Genotyping errors in a set of genotype calls considered to be accurate (the truth set), for instance, will result in an overestimation of genotyping error rates. On the other hand, consistent biases in genotype calls affecting replicates similarly, such as polymorphisms in restriction cut sites or unequal PCR amplification rates of alleles, might be difficult to infer and result in an underestimation of genotyping error rates. Finally, the assumption of strict random mating may often be violated, leading to an overestimation when inferring error rates from population samples. But despite these caveats, recalibrated genotype likelihoods are likely to reflect genotype uncertainty much more accurately, particularly for protocols with error rates as high as those reported here for RAD-seq. Indeed, we found that genotype recalibration was essential to avoid drawing inaccurate conclusions and instead recovered biologically meaningful results about the ancestry of *Populus* hybrids.

We also emphasize that if no tools accepting genotype likelihoods as input information are available for specific applications, this should not discourage users from incorporating genotyping uncertainty in the analyses. We have shown here that local ancestries were more reliably estimated by *RASPberry* (Wegmann et al., 2011), a tool requiring genotype calls, when adding the estimated genotyping error rate to the parameters of the model. But we note that given the particular lack of heterozygous genotypes in the RAD-seq data analyzed here, a model using a single per-genotype error rate as implemented in *RASPberry* was not sufficient to overcome all biases.

### Recommendations and Conclusions

Based on these results and in line with others (Cariou et al., 2016; Cooke et al., 2016; Mastretta-Yanes et al., 2015), we strongly suggest to carefully assess genotyping error rates in reduced representation sequencing experiments, and to properly account for these in downstream analyses. Here we present *Tiger*, a suite of tools to estimate genotyping error rates directly from hard genotype calls as produced by all commonly used RAD-seq analysis pipelines. Knowledge of these error rates then allows for fine-tuning the bioinformatic analysis to reduce these errors, as well as for recalibrating genotype likelihoods and hence properly accounting for genotyping errors in downstream analyses, rather than losing a large amount of information due to stringent filtering.

Among the different methods to estimate genotyping error rates, the use of population samples appears as a particularly cost-effective strategy, since no extra individuals have to be sequenced. However, not all applications include sufficiently many samples from randomly mating populations, and our simulations suggested that 20 samples are at the lower end, unless very many polymorphic sites are available. While *Tiger* can handle also multiple population samples with fewer individuals each, we expect much lower accuracy if fewer than, say, ten individuals were used per population. Also, we caution that the violation of random mating such as population sub-structure or inbreeding that causes a deficit of heterozygous genotypes directly translates into an overestimation of the heterozygous genotypes (Supplementary Figure S4).

For many studies, the preferred approach might thus be to prepare replicate libraries for a few individuals, not least because we found that just a few replicates (around ten) resulted in rather accurate estimates of genotyping errors. However, as already mentioned above, systematic biases affecting all replicates similarly may result in a slight underestimation of error rates.

Based on our simulations, the most accurate method to estimate genotyping errors is based on a truth set against which genotype calls are compared. However, this is likely also the most expensive approach. Since errors in the truth set will be wrongly interpreted as errors in the RAD-seq data, the truth set must consist of highly accurate calls. As examined here using simulations for the case of shot-gun sequencing, such a high accuracy requires considerable sequencing depth (>25x), which is likely an underestimate given the simplicity of the model used. In addition, some genotyping errors will likely prevail at low rates even in truth sets generated at very high depth, as was also reported previously (Wall et al., 2014). But given the high error rates we observed for RAD-seq data, the resulting slight overestimation of RAD-seq genotyping errors may be tolerable.

Nonetheless, and thanks to the ever dropping costs for both sequencing and library preparation, low-depth whole genome sequencing may soon become a valuable alternative to reduced representation sequencing for many applications. Since low-depth shotgun sequencing is much less prone to biases, it not only allows to interrogate a much larger fraction of the genome, but also yields accurate and precise estimates of population genetics parameters (e.g. Buerkle & Gompert, 2013; Kousathanas et al., 2017; Rustagi et al., 2017). The problem of high error rates in RAD-seq and other reduced representation libraries is thus likely transient and we expect that the field will quickly adopt new sequencing technologies that circumvent it entirely.

## Acknowledgements

Our co-author Christian Lexer passed away suddenly and prior to the final revision and submission of this manuscript. We dedicate this article to the memory of our colleague, mentor and friend Christian Lexer. We thank Kai N. Stölting, Camille Christe and Margot Paris for helpful insights and discussions, Thelma Barbará for help in the laboratory, David Frey for collecting seeds in the hybrid zone, and Santiago González Martínez for providing the Spanish samples. This work was supported by grant 31003A_149306 of the Swiss National Foundation to CL. All calculations were done at the Bioinformatic Core Facilities of the Universities of Bern and Fribourg, and computers at the University of Wyoming.

## Data Accessibility Statement

The code of Tiger is available through a git repository at https://bitbucket.org/wegmannlab/tiger. The RAD-seq data are available in the Sequence Read Archive through bioprojects PRJNA528699 and PRJNA528706. The called genotypes used to estimate genotyping errors are available at Zenodo (DOI 10.5281/zenodo.2604109 and 10.5281/zenodo.2604124).

## Author Contributions

LB and DW conceived the study; CL and DW provided funding; LB collected genetic data; LB, CAB, VL and DW performed the analyses; CAB, CL and DW supervised the study; LB and DW wrote the manuscript with input and revisions from all co-authors.

