## Supplementary Material for "Estimating and accounting for genotyping errors in RAD-seq experiments"

### Supporting Information

Luisa Bresadola[1,*], Vivian Link[1,2,*], C. Alex Buerkle[3], Christian Lexer[3], and Daniel Wegmann[1,2]

[1] *Department of Biology, University of Fribourg, Fribourg, Switzerland*
[2] *Swiss Institute of Bioinformatics, Fribourg, Switzerland*
[3] *Department of Botany, University of Wyoming, Laramie, WY, USA*
[4] *Department of Botany and Biodiversity Research, University of Vienna, Vienna, Austria*
[*] *These authors contributed equally*

**Supplementary Tables** **6**

**Supplementary Figures** **7**

### Inferring Genotyping Error Rates

#### Error rate Model

Here we describe algorithms to infer per-allele genotyping error rates from hard genotype calls $g_{il}$ obtained for individuals $i = 1, \ldots, I$ at loci $l = 1, \ldots, L$. Let $g_{il} = 0, 1, 2$ reflects the number of reference alleles and $\gamma_{il} = 0, 1, 2$ the corresponding true genotypes. Further, $\boldsymbol{g} = \{g_{11}, \ldots, g_{1L}, \ldots, g_{IL}\}$ and $\boldsymbol{\gamma} = \{\gamma_{11}, \ldots, \gamma_{1L}, \ldots, \gamma_{IL}\}$.

We assume a simple error model with the per allele genotyping error rates $\boldsymbol{\epsilon} = \{\epsilon_0, \epsilon_1\}$ such that

$$\mathbb{P}(g|\gamma, \epsilon_0, \epsilon_1) = \begin{cases} (1-\epsilon_0)^2 & \text{if } g = \gamma; \gamma = 0, 2 \\ \epsilon_1^2 + (1-\epsilon_1)^2 & \text{if } g = \gamma = 1 \\ \epsilon_0^2 & \text{if } g = 0; \gamma = 2 \text{ or } g = 2; \gamma = 0 \\ 2\epsilon_0(1-\epsilon_0) & \text{if } g = 1; \gamma = 0, 2 \\ \epsilon(1-\epsilon_1) & \text{if } g = 0, 2; \gamma = 1 \end{cases} . \quad (1)$$

Often, not all genotype calls are affected by the same genotyping error rates. For instance, genotyping error rates might vary between calls obtained with different sequencing depth or between samples of different batches such as from different libraries or sequencing experiments. We account for this by stratifying the error rates by different sets $s = 1, \ldots, S$ of genotypes and denote the corresponding error rates by $\epsilon_0(s)$ and $\epsilon_1(s)$, respectively. We further denote by $s_{il}$ the set to which genotype call $g_{il}$ belongs. Finally, $\boldsymbol{\epsilon}_0 = \{\epsilon_0(1), \ldots, \epsilon_0(S)\}$, $\boldsymbol{\epsilon}_1 = \{\epsilon_1(1), \ldots, \epsilon_1(S)\}$ and $\boldsymbol{\epsilon} = \{\boldsymbol{\epsilon}_0, \boldsymbol{\epsilon}_1\}$.

#### Inferring Genotyping Error Rates by Comparing to a Truth Set

We first describe an algorithm to infer per-allele genotyping error rates by comparing calls to a truth set, i.e. the case in which all $\gamma_{il}$ are known. Under the assumption that genotyping errors are independent between loci and individuals, the likelihood of the full data is given by

$$\mathbb{P}(\boldsymbol{g}|\boldsymbol{\gamma}, \epsilon) = \prod_{i=1}^{I}\prod_{l=1}^{L} \mathbb{P}(g_{il}|\gamma_{il}, \boldsymbol{\epsilon})$$

where $\mathbb{P}(g_{il}|\gamma_{il}, \boldsymbol{\epsilon}) = \mathbb{P}(g_{il}|\gamma_{il}, \epsilon_0(s_{il}), \epsilon_1(s_{il}))$ according to (1).

In order to obtain a maximum likelihood (ML) estimate of $\epsilon$, we numerically maximize the likelihood function by finding $\epsilon$ such that

$$\frac{\partial}{\partial \epsilon} \log \mathbb{P}(\boldsymbol{g}|\boldsymbol{\gamma}, \boldsymbol{\epsilon}) = 0.$$

Consider a table with entries $n_{g,\gamma}(s)$ reflecting the counts of observing genotype $g$ at a site with true genotype $\gamma$ for all calls of set $s$. Given this table, the log-likelihood function can be calculated as

$$\begin{aligned} \log \mathbb{P}(\boldsymbol{n}|\boldsymbol{\epsilon}) &= \sum_{g=0}^{2}\sum_{\gamma=0}^{2} n_{g,\gamma}(s) \log \mathbb{P}(g|\gamma, \epsilon) \\ &= \sum_{s} a_0(s) \log \epsilon_0(s) + b_0(s) \log(1-\epsilon_0(s)) + \ldots \\ &\quad \ldots + a_1(s) \log \epsilon_1(s) + b_1(s) \log(1-\epsilon_1(s)) + c(s) \log(\epsilon_1^2(s) + (1-\epsilon_1(s))^2) + C \end{aligned}$$

where $C$ is a constant independent of $\boldsymbol{\epsilon}$ and

$$\begin{aligned} a_0(s) &= 2n_{0,2}(s) + n_{1,0}(s) + n_{1,2}(s) + 2n_{2,0}(s) \\ a_1(s) &= n_{0,1}(s) + n_{2,1}(s) \\ b_0(s) &= 2n_{0,0}(s) + n_{1,0}(s) + n_{1,2}(s) + 2n_{2,2}(s) \\ b_1 &= n_{0,1} + n_{2,1} \\ c_0 &= n_{1,1} \end{aligned}$$

From this we get for $\epsilon_0(s)$

$$\frac{\partial}{\partial \epsilon_0(s)} \log \mathbb{P}(\boldsymbol{n}|\boldsymbol{\epsilon}) = \frac{a}{\epsilon_0(s)} - \frac{b}{1-\epsilon_0(s)} = 0,$$

and the MLE estimate

$$\hat{\epsilon_0(s)} = \frac{a_0(s)}{a_0(s) + b_0(s)}.$$

For $\epsilon_1(s)$ we get

$$\begin{aligned} \frac{\partial}{\partial \epsilon_1(s)} \log \mathbb{P}(\boldsymbol{n}|\boldsymbol{\epsilon}) &= -2[a_1(s) + b_1(s) + 2c(s)]\epsilon_1(s)^3 + 2[2a_1(s) + b_1(s) + 3c(s)]\epsilon_1(s)^2 - \dots \\ &\quad \dots - [3a_1(s) + b_1(s) + 2c(s)]\epsilon_1(s) + a_1(s), \end{aligned}$$

which we set to zero using a numerical search algorithm to obtain the MLE estimate $\hat{\epsilon_1(s)}$.

#### Inferring Genotyping Error Rates from Sample Replicates

We next develop an algorithm to infer genotyping error rates from individuals for which genotypes were estimated from multiple, independent replicate data sets. Consider inferred genotypes $\boldsymbol{g}_i = \{\boldsymbol{g}_i^{(1)}, \dots, \boldsymbol{g}_i^{(r_i)}\}$ for multiple individuals $i = 1, \dots, I$, where $\boldsymbol{g}_i^{(j)} = \{g_{i1}^{(j)}, \dots, g_{iL}^{(j)}\}$ indicates the inferred genotypes of individual $i$ at sites $l = 1 \dots, L$ from the $j$th replicate. The likelihood of the full data is then given by

$$\mathbb{P}(\boldsymbol{g}|\boldsymbol{\epsilon}) = \prod_{i=1}^{I} \prod_{l=1}^{L} \sum_{g=0}^{2} \mathbb{P}(\gamma_{il} = g) \prod_{j=1}^{r_i} \mathbb{P}(g_{il}^{(j)} | \gamma_{il} = g, \epsilon_0(s_{il}^{(j)}), \epsilon_1(s_{il}^{(j)})), \tag{2}$$

where $\mathbb{P}(\gamma_{il} = g) = f_{ig}$ is the frequency of genotype $g$ among all sites typed in individual $i$, $s_{il}^{(j)}$ the set to which call $g_{il}^{(j)}$ belongs, and $\mathbb{P}(g_{il}^{(j)} | \gamma_{il} = g, \epsilon_0(s_{il}^{(j)}), \epsilon_1(s_{il}^{(j)}))$ is given by (1).

In order to find the MLE estimates of $\epsilon$ and $\boldsymbol{f} = \{f_i, \dots, f_I\}$, $f_i = \{f_{i0}, f_{i1}, f_{i2}\}$ we will employ an EM algorithm.

**E-step.** The expected complete data likelihood is given by

$$\begin{aligned} Q(\boldsymbol{\epsilon}, \boldsymbol{f}; \boldsymbol{\epsilon}', \boldsymbol{f}') &= \mathbb{E}[\ell_c(\boldsymbol{\epsilon}, \boldsymbol{f}) | \boldsymbol{g}, \boldsymbol{\epsilon}', \boldsymbol{f}'] \\ &= \sum_{i=1}^{I} \sum_{l=1}^{L} \sum_{g=0}^{2} \mathbb{P}\left(\gamma_{il} = g | \boldsymbol{g}_{il}, \epsilon_0'(s_{il}^{(j)}), \epsilon_1'(s_{il}^{(j)}), \boldsymbol{f}'\right) \left[\log f_{ig} + \sum_{j=1}^{r_i} \log \mathbb{P}\left(g_{il}^{(j)} | \gamma_{il} = g, \epsilon_0'(s_{il}^{(j)}), \epsilon_1'(s_{il}^{(j)})\right)\right], \end{aligned}$$

where we used $\boldsymbol{g}_{il} = \{g_{il}^{(1)}, \dots, g_{il}^{(r_i)}\}$ and we have, according to Bayes formula,

$$\mathbb{P}\left(\gamma_{il} = g | \boldsymbol{g}_{il}, \epsilon_0'(s_{il}^{(j)}), \epsilon_1'(s_{il}^{(j)}), \boldsymbol{f}'\right) = \frac{f_{ig}' \prod_{j=1}^{r_i} \mathbb{P}\left(g_{il}^{(j)} | \gamma_{il} = g, \epsilon_0'(s_{il}^{(j)}), \epsilon_1'(s_{il}^{(j)})\right)}{\sum_{h=0}^{2} f_{ih}' \prod_{j=1}^{r_i} \mathbb{P}\left(g_{il}^{(j)} | \gamma_{il} = h, \epsilon_0'(s_{il}^{(j)}), \epsilon_1'(s_{il}^{(j)})\right)}.$$

**M-step.** We have to maximize $Q$ subject to the constraints

$$\sum_{g=0}^{2} f_{ig} = f_{i0} + f_{i1} + f_{i2} = 1 \tag{3}$$

for all individuals $i = 1, \dots, I$, we form the Lagrangian

$$\mathcal{L}(\boldsymbol{\epsilon}, \boldsymbol{f}, \boldsymbol{\mu}) = Q - \sum_{i=0}^{I} \mu_i \left( \sum_{g=0}^{2} f_{ig} - 1 \right), \tag{4}$$

where $\boldsymbol{\mu} = \{\mu_1, \dots, \mu_I\}$ is the vector of Lagrangian multipliers. All $\boldsymbol{f}$ can be maximized analytically. We get the following derivatives of the Lagrangian:

$$\begin{aligned}
\frac{\partial}{\partial \mu_i}\mathcal{L} &= \sum_{g=0}^{2} f_{ig} - 1 = 0 \\
\frac{\partial}{\partial f_{ig}}\mathcal{L} &= \frac{1}{f_{ig}} \sum_{l=1}^{L} \mathbb{P}\left(\gamma_{il} = g | \boldsymbol{g}_{il}, \epsilon'_0(s_{il}^{(j)}), \epsilon'_1(s_{il}^{(j)}), \boldsymbol{f}'\right) - \mu_i = 0.
\end{aligned}$$

From these we get

$$f_{ig} = \frac{1}{L} \sum_{l=1}^{L} \mathbb{P}\left(\gamma_{il} = g | \boldsymbol{g}_{il}, \epsilon'_0(s_{il}^{(j)}), \epsilon'_1(s_{il}^{(j)}), \boldsymbol{f}'\right).$$

In contrast, we will need to maximize $Q$ with respect to each $\epsilon_0(s)$ and $\epsilon_1(s)$ numerically. For this, we need to maximize

$$\begin{aligned}
Q_{\epsilon_0(s)} &= \sum_{i=1}^{\mathcal{I}} \sum_{l=1}^{L} \sum_{g \in \{0,2\}} I(s_{il}^{(j)} = s) \mathbb{P}(\gamma_{il} = g | \boldsymbol{g}_{il}, \epsilon'_0(s), \boldsymbol{f}') \sum_{j=1}^{r_i} \log \mathbb{P}(g_{il}^{(j)} | \gamma_{il} = g, \epsilon_0(s)), \\
Q_{\epsilon_1(s)} &= \sum_{i=1}^{\mathcal{I}} \sum_{l=1}^{L} I(s_{il}^{(j)} = s) \mathbb{P}(\gamma_{il} = 1 | \boldsymbol{g}_{il}, \epsilon_1(s), \boldsymbol{f}') \sum_{j=1}^{r_i} \log \mathbb{P}(g_{il}^{(j)} | \gamma_{il} = 1, \epsilon_1(s)),
\end{aligned}$$

where the sums run only over the genotypes of set $s$, as given by the indicator function

$$\mathcal{I}(s_{il}^{(j)} = s) = \begin{cases} 1 & \text{if } s_{il}^{(j)} = s, \\ 0 & \text{otherwise.} \end{cases}$$

#### Inferring Genotyping Error Rates from Population Samples

We finally develop an algorithm to infer genotyping error rates from multiple samples from each of multiple populations. Consider observed genotypes $g_{pil}$ from individual $i = 1 \dots, n_p$ from population $p = 1, \dots, P$ at locus $l = 1, \dots, L$. Let us further denote by $\boldsymbol{g}_{pi} = \{g_{pi1}, \dots, g_{piL}\}$ and by $\boldsymbol{g}_p = \{\boldsymbol{g}_{p1}, \dots, \boldsymbol{g}_{pn_p}\}$. The likelihood of the full data given per allele genotyping error rates $\boldsymbol{\epsilon} = \{\epsilon_1, \dots, \epsilon_S\}$ for different sets $s = 1, \dots, S$ of genotype calls is then given by

$$\mathbb{P}(\boldsymbol{g}|\boldsymbol{\epsilon}) = \prod_{p=1}^{P} \prod_{i=1}^{n_p} \prod_{l=1}^{L} \sum_{g=0}^{2} \mathbb{P}(\gamma_{pil} = g | f_{pl}) \mathbb{P}\left(g_{pil} | \gamma_{pil} = g, \epsilon_0(s_{pil}), \epsilon_1(s_{pil})\right), \tag{5}$$

where $\boldsymbol{g} = \{\boldsymbol{g}_1, \dots, \boldsymbol{g}_P\}$, $f_{pl}$ is the frequency in population $p$ of one of the two alleles at locus $l$, and under the assumption of Hardy-Weinberg equailibrium in each population,

$$\mathbb{P}(g_{pil}|f_{pl}) = \binom{2}{g_{pil}} f_{pl}^{g_{pil}} (1 - f_{pl})^{2 - g_{pil}} = \begin{cases} (1 - f_{pl})^2 & \text{if } g_{pil} = 0, \\ 2f(1 - f_{pl}) & \text{if } g_{pil} = 1, \\ f_{pl}^2 & \text{if } g_{pil} = 2. \end{cases} \tag{6}$$

Finally, $s_{pil}$ is the set of call $g_{pil}$ and $\mathbb{P}\left(g_{il}^{(j)} | \gamma_{il}, \epsilon_0(s), \epsilon_1(s)\right)$ is given by (1).

#### Maximum Likelihood Inference

In order to find the MLE estimates of $\boldsymbol{\epsilon}$ and $\boldsymbol{f} = \{\boldsymbol{f}_1, \ldots, \boldsymbol{f}_P\}$, $\boldsymbol{f}_p = \{f_{p1}, \ldots, f_{pL}\}$ we will employ an EM algorithm.

**E-step.** The expected complete data likelihood is given by

$$\begin{aligned} Q(\boldsymbol{\epsilon}, \boldsymbol{f}; \boldsymbol{\epsilon}', \boldsymbol{f}') &= \mathbb{E}[\ell_c(\boldsymbol{\epsilon}, \boldsymbol{f})|\boldsymbol{g}, \boldsymbol{\epsilon}', \boldsymbol{f}'] \\ &= \sum_{p=1}^{P}\sum_{i=1}^{n_p}\sum_{l=1}^{L}\sum_{g=0}^{2} \mathbb{P}\left(\gamma_{pil} = g|\boldsymbol{g}_{pil}, \epsilon_0'(s_{pil}), \epsilon_1'(s_{pil}), f_{plg}'\right) \Big[\log f_{plg} + \log \mathbb{P}\left(g_{pil}|\gamma_{pil} = g, \epsilon_0(s_{pil}), \epsilon_1(s_{pil})\right)\Big], \end{aligned}$$

According to Bayes formula,

$$\mathbb{P}\left(\gamma_{pil} = g|\boldsymbol{g}_{pil}, \epsilon_0(s_{pil}), \epsilon_1(s_{pil}), f_{plg}'\right) = \frac{\mathbb{P}\left(g_{pil}|\gamma_{pil} = g, \epsilon_0(s_{pil}), \epsilon_1(s_{pil})\right) f_{plg}'}{\sum_{h=0}^{2} \mathbb{P}\left(g_{pil}|\gamma_{pil} = h, \epsilon_0(s_{pil}), \epsilon_1(s_{pil})\right) f_{plh}'}.$$

**M-step.** We have to maximize $Q$ subject to the constraints

$$\sum_{g=0}^{2} f_{plg} = f_{pl0} + f_{pl1} + f_{pl2} = 1$$

for all populations $p = 1, \ldots, P$, we form the Lagrangian

$$\mathcal{L}(\boldsymbol{\epsilon}, \boldsymbol{f}, \boldsymbol{\mu}) = Q - \sum_{p=1}^{P} \mu_p \left(\sum_{g=0}^{2} f_{plg} - 1\right),$$

where $\boldsymbol{\mu} = \{\mu_1, \ldots, \mu_P\}$ is the vector of Lagrangian multipliers. All $\boldsymbol{f}$ can be maximized analytically. We get the following derivatives of the Lagrangian:

$$\begin{aligned} \frac{\partial}{\partial \mu_p}\mathcal{L} &= \sum_{g=0}^{2} f_{plg} - 1 = 0 \\ \frac{\partial}{\partial f_{plg}}\mathcal{L} &= \frac{1}{f_{plg}} \sum_{i=1}^{n_p} \mathbb{P}(\gamma_{pil} = g|\boldsymbol{g}_{pil}, \epsilon_0'(s_{pil}), \epsilon_1'(s_{pil}), f_{plg}') - \mu_p = 0. \end{aligned}$$

From these we get

$$f_{plg} = \frac{1}{n_p} \sum_{i=1}^{n_p} \mathbb{P}(\gamma_{pil} = g|\boldsymbol{g}_{pil}, \epsilon_0'(s_{pil}), \epsilon_1'(s_{pil}), f_{plg}').$$

In contrast, we will need to maximize $Q$ with respect to each $\epsilon_s$ numerically. For this, we need to maximize

$$Q_{\epsilon_s} = \sum_{p=1}^{P}\sum_{i=1}^{n_p}\sum_{l=1}^{L} \mathcal{I}(s_{pil} = s) \sum_{g=0}^{2} \mathbb{P}\left(\gamma_{pil} = g|\boldsymbol{g}_{pil}, \epsilon_0'(s_{pil}), \epsilon_1'(s_{pil}), f_{plg}'\right) \Big[\log f_{plg} + \log \mathbb{P}\left(g_{pil}|\gamma_{pil} = g, \epsilon_0(s_{pil}), \epsilon_1(s_{pil})\right)\Big],$$

where the sums run only over the genotypes of set $s$, as given by the indicator function

$$\mathcal{I}(s_{pil} = c) = \begin{cases} 1 & \text{if } s_{pil} = s, \\ 0 & \text{otherwise.} \end{cases}$$

#### Bayesian Inference

The MLE estimator introduced above results in an underestimation of the error rates because the allele frequencies are estimated too close to the data, i.e. the MLE of the allele frequencies are biased. Importantly, this problem only disappears when using excessively many samples, but not when using more loci since an allele frequency must be estimated for every locus. To obtain meaningful estimates also for reasonable sample sizes, we develop a Bayesian inference method for the posterior distribution that integrates over the uncertainty of allele frequencies. This results in an unbiased estimation if sufficiently many loci are used (see Figure 1 in the main text).

$$\mathbb{P}(\boldsymbol{g}|\boldsymbol{\epsilon},\boldsymbol{f}) = \frac{\mathbb{P}(\boldsymbol{g}|\boldsymbol{\epsilon},\boldsymbol{f})\mathbb{P}(\boldsymbol{\epsilon})\mathbb{P}(\boldsymbol{f})}{\mathbb{P}(\boldsymbol{\epsilon},\boldsymbol{f})}, \tag{7}$$

assuming an exponential prior on $\boldsymbol{\epsilon}$

$$\mathbb{P}(\boldsymbol{\epsilon}) = \prod_{s=1}^{C}[\mathbb{P}(\epsilon_0(s))\mathbb{P}(\epsilon_1(s))]; \quad \epsilon_0(s), \epsilon_1(s) \sim \text{Exponential}(\lambda)$$

and the uninformative Jeffrey's prior on $\mathbb{P}(\boldsymbol{f})$

$$\mathbb{P}(\boldsymbol{f}) = \prod_{p=1}^{P}\prod_{l=1}^{L}\mathbb{P}(f_{pl}); \quad \mathbb{P}(f_{pl}) \sim \text{Beta}(0.5, 0.5)$$

Since $\mathbb{P}(\boldsymbol{\epsilon},\boldsymbol{f})$ can not be evaluated analytically, we resort to an MCMC algorithm to generate samples from the posterior distribution $\mathbb{P}(\boldsymbol{g}|\boldsymbol{\epsilon},\boldsymbol{f})$.

### Supplementary Tables

**Table S1:** Samples included in the RAD vs GBS comparison. The RAD data is available through the SRA Biproject PRJNA528699. Compared genotypes are available at Zenodo through DOI 10.5281/zenodo.2604109.

| | | | | | | | | | |
|---|---|---|---|---|---|---|---|---|---|
| F008_01 | F008_15 | F009_12 | F021_06 | F026_06 | F030_03 | F032_03 | F033_05 | F039_05 | I373_05 |
| F008_02 | F008_16 | F009_13 | F021_07 | F026_07 | F030_04 | F032_04 | F033_06 | F039_06 | I373_06 |
| F008_03 | F008_17 | F020_01 | F022_01 | F026_08 | F030_05 | F032_05 | F033_07 | F039_07 | I373_07 |
| F008_04 | F009_01 | F020_02 | F022_02 | F026_09 | F030_06 | F032_06 | F036_01 | I345_01 | I396_01 |
| F008_05 | F009_02 | F020_03 | F022_03 | F026_10 | F030_07 | F032_08 | F036_02 | I345_02 | I396_02 |
| F008_06 | F009_03 | F020_04 | F022_04 | F026_11 | F031_01 | F032_09 | F036_03 | I345_03 | I396_03 |
| F008_07 | F009_04 | F020_05 | F022_05 | F026_12 | F031_02 | F032_10 | F036_04 | I345_04 | I396_04 |
| F008_08 | F009_05 | F020_06 | F022_06 | F026_13 | F031_03 | F032_11 | F036_05 | I345_05 | I396_05 |
| F008_09 | F009_06 | F020_07 | F022_07 | F026_14 | F031_04 | F032_12 | F036_06 | I345_06 | I396_06 |
| F008_10 | F009_07 | F021_01 | F026_01 | F026_15 | F031_05 | F032_13 | F036_07 | I345_07 | I396_07 |
| F008_11 | F009_08 | F021_02 | F026_02 | F026_16 | F031_06 | F033_01 | F039_01 | I373_01 | |
| F008_12 | F009_09 | F021_03 | F026_03 | F026_17 | F031_07 | F033_02 | F039_02 | I373_02 | |
| F008_13 | F009_10 | F021_04 | F026_04 | F030_01 | F032_01 | F033_03 | F039_03 | I373_03 | |
| F008_14 | F009_11 | F021_05 | F026_05 | F030_02 | F032_02 | F033_04 | F039_04 | I373_04 | |

### Supplementary Figures

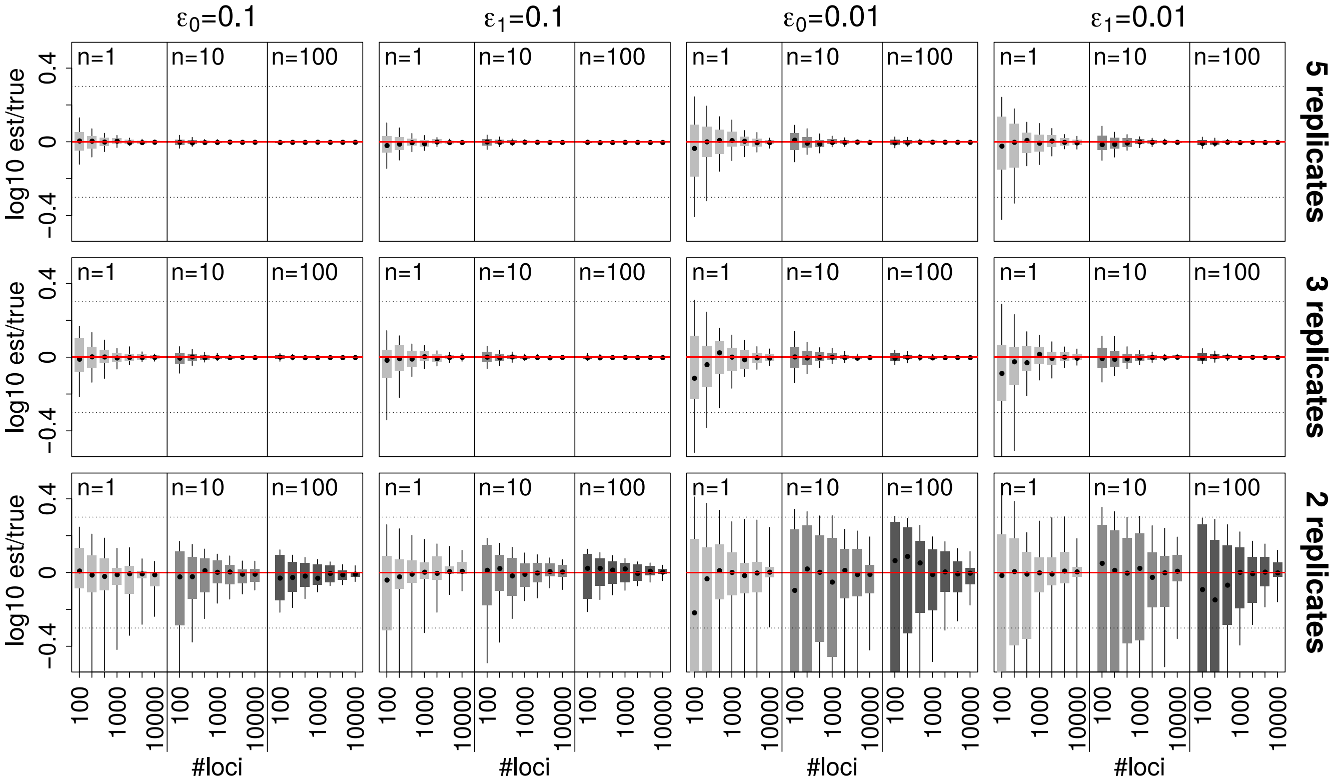

**Figure S1:** Accuracy of error rate estimates from two (bottom row), three (middle row) and five (top row) replicates per samples. Shown are the estimated error rates relative to the true error rates (red line) of 100 replicates for different samples sizes n (shown on top), the two error rates 0.1 and 0.01 (first and last two columns, respectively) and different numbers of unlinked loci. The horizontal dashed lines indicate Q2, the interval within which an estimate is less than a factor of two away from the true value.

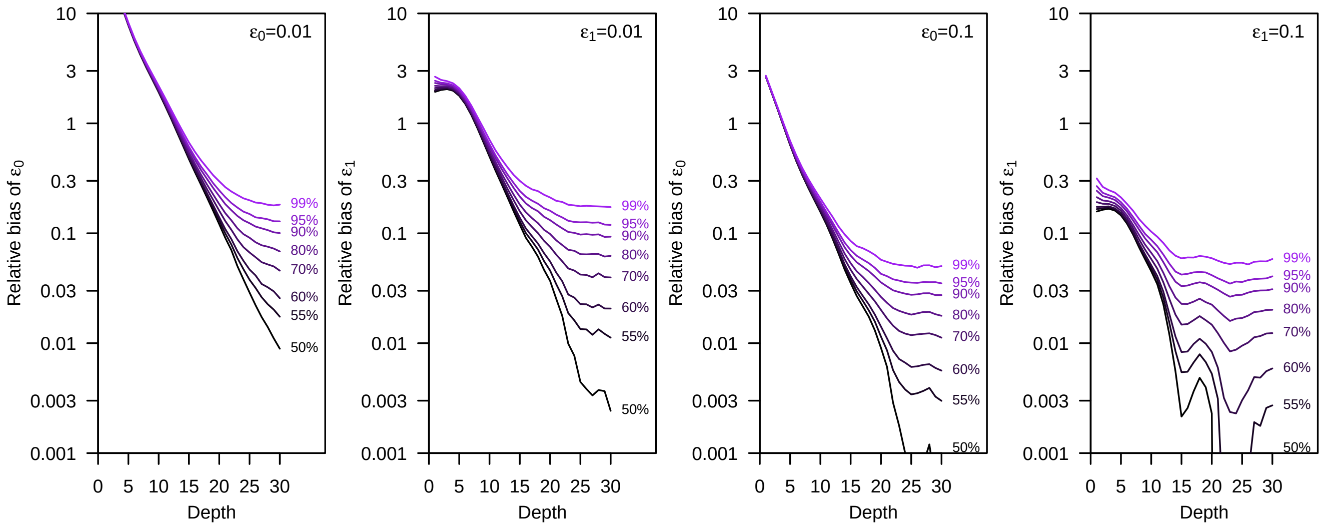

**Figure S2:** Distribution of the relative bias when inferring error rates from a truth set generated with next-generation sequencing data with a specific mean depth. Each line indicates a quantile q such that a smaller bias was observed in a fraction q of $10^6$ simulations of $10^4$ homozygous and $10^4$ heterozygous loci. The left two panels are for the case $\epsilon_0 = \epsilon_1 = 0.01$ and the right two panels for $epsilon_0 = \epsilon_1 = 0.1$.

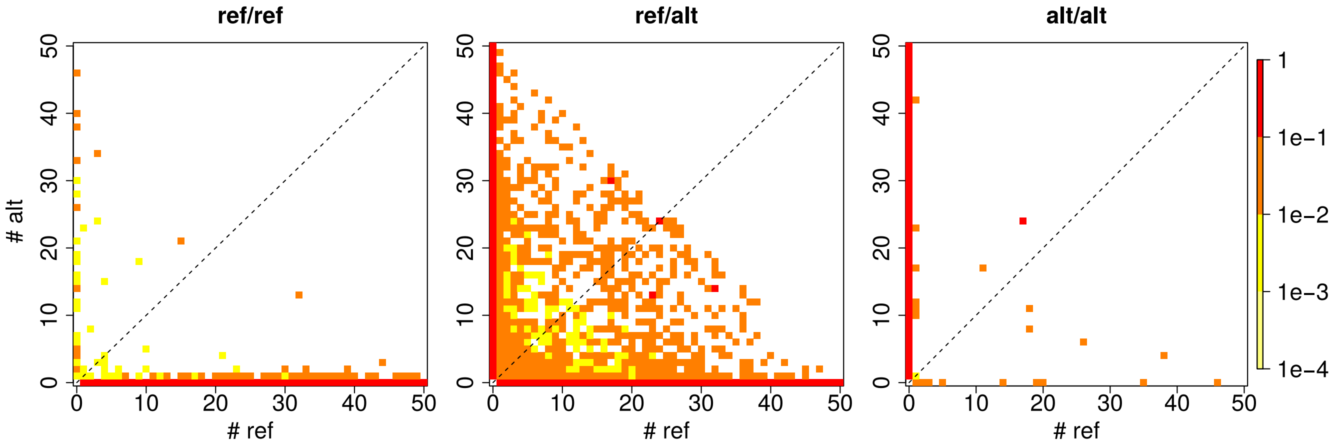

**Figure S3:** Distribution of the allelic depth observed in the RAD-seq data for genotypes found to be homozygous reference (left, 8,701 genotypes), heterozygous (middle, 6,022 genotypes) and homozygous alternative (right, 1,914 genotypes) genotypes in the GBS truth set exploiting familial relationships. Allelic depth distributions are normalized per total depth for plotting (i.e. they sum to 1 per top-left to lower-right diagonal).

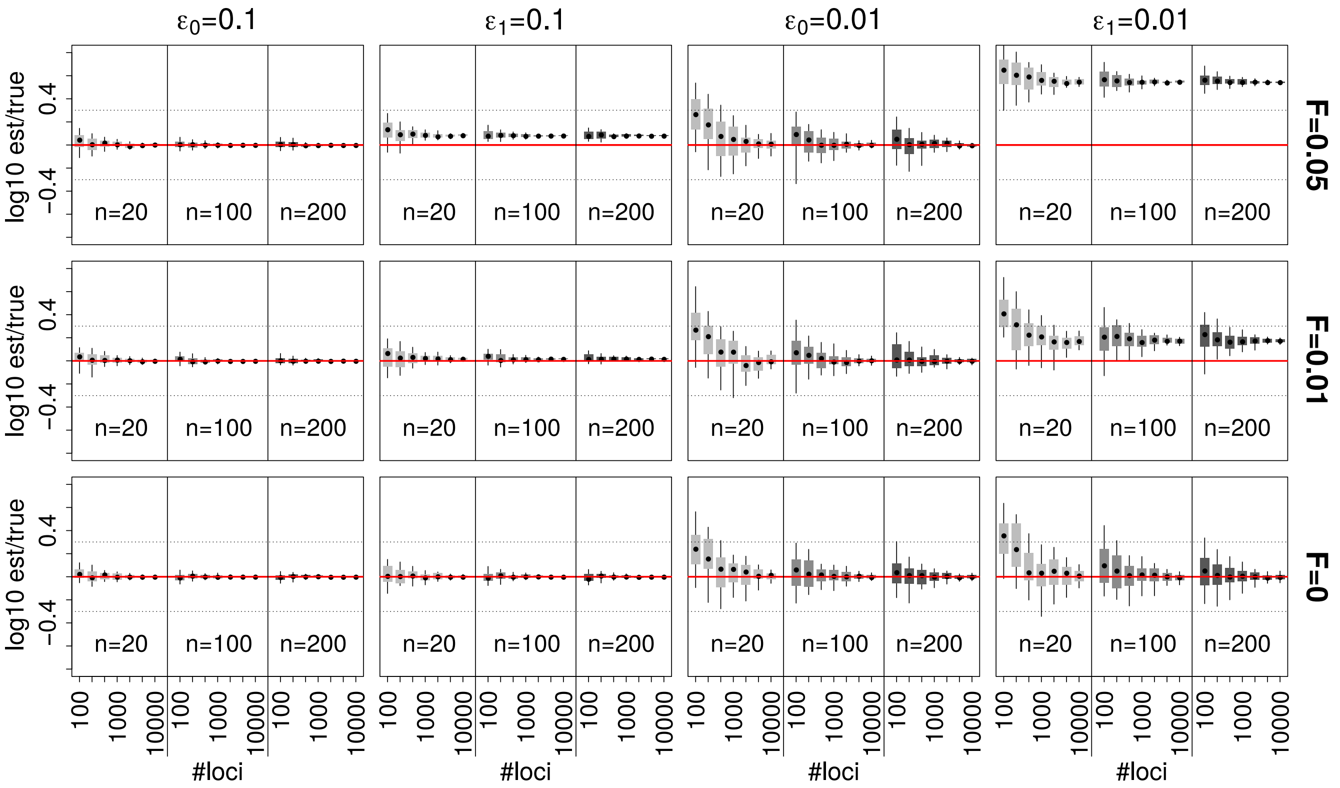

**Figure S4:** Accuracy of error rate estimates under the population sample model if data deviates from Hardy-Weinberg proportion as quantified by the inbreeding coefficient $F$. Shown are the estimated error rates relative to the true error rates (red line) of 100 replicates for different samples sizes n (shown on top), the two error rates 0.1 and 0.01 (first and last two columns, respectively) and different numbers of unlinked loci. The horizontal dashed lines indicate Q2, the interval within which an estimate is less than a factor of two away from the true value.
